# Expression changes in immune and epigenetic gene pathways associated with nutritional metabolites in maternal blood from pregnancies resulting in autism and atypical neurodevelopment

**DOI:** 10.1101/2020.05.14.096586

**Authors:** Yihui Zhu, Charles E. Mordaunt, Blythe P Durbin-Johnson, Marie A Caudill, Olga V. Malysheva, Joshua W. Miller, Ralph Green, S. Jill James, Stepan B. Melnyk, M. Daniele Fallin, Irva Hertz-Picciotto, Rebecca J. Schmidt, Janine M. LaSalle

## Abstract

**Background:** The prenatal period is a critical window to study factors involved in the development of autism spectrum disorder (ASD). Environmental factors, especially *in utero* nutrition, can interact with genetic risk for ASD, but how specific prenatal nutrients in mothers of children later diagnosed with ASD or non-typical development (Non-TD) associate with gestational gene expression is poorly understood. Maternal blood collected prospectively during pregnancy provides a new opportunity to gain insights into nutrition, particularly one-carbon metabolites, on gene pathways and neurodevelopment.

**Methods:** Genome-wide transcriptomes were measured using microarrays in 300 maternal blood samples from all three trimesters in the Markers of Autism Risk in Babies - Learning Early Signs (MARBLES) study. Sixteen different one-carbon metabolites, including folic acid, betaine, 5’-methyltretrahydrofolate (5-MeTHF), and dimethylglycine (DMG) were measured. Differential expression analysis and weighted gene correlation network analysis (WGCNA) were used to compare gene expression between children later diagnosed as typical development (TD), Non-TD and ASD, and to nutrient metabolites.

**Results:** Using differential gene expression analysis, six transcripts associated with four genes (*TGR-AS1, SQSTM1, HLA-C* and *RFESD*) showed genome-wide significance (FDR *q* < 0.05) with child outcomes. Genes nominally differentially expressed compared to TD specifically in ASD, but not Non-TD, significantly overlapped with seven high confidence ASD genes. 218 transcripts in common to ASD and Non-TD differential expression compared to TD were significantly enriched for functions in immune response to interferon-gamma, apoptosis, and metal ion transport. WGCNA identified co-expressed gene modules significantly correlated with 5-MeTHF, folic acid, DMG, and betaine. A module enriched in DNA methylation functions showed a protective association with folic acid/5-MeTHF concentrations and ASD risk. Independent of child outcome, maternal plasma betaine and DMG concentrations associated with a block of co-expressed genes enriched for adaptive immune, histone modification, and RNA processing functions.

**Limitations:** Blood contains a heterogeneous mixture of cell types, and many WGCNA modules correlated with cell type and/or nutrient concentrations, but not child outcome. Gestational age correlated with some co-expressed gene modules in addition to nutrients.

**Conclusions:** These results support the premise that the prenatal maternal blood transcriptome is a sensitive indicator of gestational nutrition and children’s later neurodevelopmental outcomes.

## Background

Autism spectrum disorder (ASD) is a group of neurodevelopmental disorders characterized by persistent impairment in social interactions, communication, restricted interests or repetitive behaviors, and sensory sensitivities [1]. Current data show that one in every 59 children in the United States has ASD [1]. One major component of ASD risk is genetic heritability, based on studies of twins, siblings, and other family members [2–4]. Common genetic variants each having small effects dominate most ASD risk compared with rare gene variants with large effects [5]. Large genome-wide association studies (GWAS) support the role of common genetic variants in ASD with remaining challenges in ASD complexity and heterogeneity [6–8]. Mutations in single genes can only explain less than 1% of ASD cases [9,10].

Accumulating lines of evidence suggest that ASD risk arises from both genetic and environmental risk factors. *In utero* maternal exposures can contribute as ASD risk factors, including air pollution, fever, asthma, and nutrition, especially nutrients involved in the one-carbon metabolic pathway [11–14]. Other studies suggest that one-carbon metabolism is implicated in gene-environment interactions in ASD [15,16]. Maternal prenatal nutritional supplements containing folic acid and additional B vitamins that play a role in one-carbon metabolism are associated with an estimated 40% ASD risk reduction [13,17,18].

Gene expression levels are also influenced by both genetic and environmental factors, especially by *in utero* maternal nutrition [19,20]. Maternal peripheral blood therefore offers a unique window to study transcriptome alterations during pregnancy that may reflect altered fetal development associated with nutrition [21,22]. Numerous environmental factors during pregnancy can alter gene expression levels [23,24]. Recent studies of schizophrenia demonstrated a significant interaction of genetic risk with maternal perinatal environmental factors that affected the transcriptome [25,26]. Postmortem brain gene expression studies showed gene co-expression was enriched at immune response and neuronal development in ASD [27,28]. Other studies using child peripheral blood and cord blood showed differential gene expression in ASD was enriched for immune and inflammatory processes [29–31].

While numerous studies have investigated specific genes or pathways in children with ASD, none have focused on the maternal transcriptome during pregnancy. Further, most previous ASD transcriptome studies used data from specimens collected postmortem or after childbirth, as opposed to prospective studies to help understand potential etiologic changes that occur before behavior symptoms. Other large epidemiology studies examined environmental effects in ASD, but how the environment influences alterations at the molecular level remains to be understood. The goal of this study was to examine maternal prenatal gene expression profiles associated with both maternal serum one-carbon metabolites and the child’s later autism diagnosis to shed light on molecular changes during pregnancy.

## Materials and Methods

### MARBLES study design

The MARBLES study recruited mothers in Northern California with at least one child with ASD who were pregnant or planning another pregnancy. A previous publication detailed the study design of MARBLES [31,32]. In order to enroll into MARBLES, all five following criteria needed to be met: (1) the fetus of interest has one or more first or second degree relatives with ASD; (2) mother is 18 years or older; (3) mother is pregnant or able to become pregnant; (4) mother is able to speak, read, and understand English and plans to raise the child with English spoken at home; (5) mother lives within 2.5-hour drive distance from Davis/Sacramento, California region. Due to a shared genetic background, the next child has higher risk for ASD. Demographic, diet, environmental, and medical information were collected by telephone interviews or questionnaires through the pregnancy. Infants received standardized neurodevelopmental assessments from 6 months until 3 years old [32]. At 3 years old, the child was assessed clinically using the gold standard Autism Diagnostic Observation Schedule (ADOS) [33], the Autism Diagnostic Interview – Revised (ADI-R) [34], and the Mullen Scales of Early Learning (MSEL) [35]. Based on a previously published algorithm using ADOS and MSEL scores [18,31], participants were classified into three outcome groups including ASD, TD, and Non-TD [36,37]. Children with ASD had scores over the ADOS cutoff and fit ASD DSM-5 criteria. Children with Non-TD outcomes were defined as children with low MSEL scores (two or more MSEL subscales with more than 1.5 standard deviations (SD) below averages or at least one MSEL subscale more than 2 SD below average) and elevated ADOS scores.

Children with TD outcome had all MSEL scores within 2.0 SD and no more than one MSEL subscale that is 1.5 SD below the normative mean and scores on the ADOS at least three points lower than the ASD cutoff.

### RNA isolation and expression microarray

Maternal peripheral blood was collected at study visits during all three trimesters of pregnancy in PAXgene Blood RNA tubes with the RNA stabilization reagent (BD Biosciences) and stored frozen at −80°C. The first timepoint was used for mothers who had multiple blood draws (n = 12) during pregnancy. RNA was isolated using the PAXgene Blood RNA Kit (Qiagen) according to the manufacturer’s instructions. Total RNA was converted to cDNA and biotin labeled. Expression was measured using Human Gene 2.0 Affymetrix microarray chips by the John Hopkins Sequencing and Microarray core following washing, staining, and scanning procedures based on manufacture’s protocol.

### Data preprocessing and normalization

Robust Multi-Chip Average (RMA) [38–40] from the oligo R package was used for normalization of Affymetrix CEL files. For quality control, we used the oligo and ArrayQualityMetrics R packages [41,42]. No samples were identified as outliers by principal component analysis, the Kolmogorov-Smirnov test, or distance to other arrays. Probes were mapped at the transcript level using the pd.hugene.2.0.st R package, and those annotated to genes (36,459) were used in subsequent analyses.

### One-carbon nutrient metabolite measurements

Serum and plasma samples from the same blood draw as specimens used for RNA expression analysis were used to measure one-carbon and nutrient metabolites. S-adenosylmethionine (SAM) and S-adenosylhomocysteine (SAH), adenosine, homocysteine, cystine, cysteine, glutathione (GSH) and glutathione disulfide (GSSG) were measured in the James’ laboratory at the Arkansas Children’s Research Institute using HPLC with electrochemical detection as previously described [43,44]. Serum pyridoxal phosphate (PLP), the biologically active form of vitamin B6 (Vit B6), was measured by HPLC using fluorescence detection in the Green-Miller laboratory at the UC Davis Medical Center (interassay CV = 4.8%) [45]. Total serum vitamin B12 (Vit B12) was measured using automated chemiluminescence in the CLIA-approved Medicine Clinical Laboratories at UC Davis Medical Center (inter-assay CV = 6.2%). Plasma choline, betaine and dimethylglycine (DMG) were measured using LC-MS/MS stable isotope dilution methods in the Caudill laboratory [46,47] with modifications to include measurements of trimethylamine N-oxide (TMAO) and methionine [48,49]. Intra-assay and inter-assay CVs of the in-house controls were 3.0% and 3.6% for choline; 1.5% and 1.7% for betaine, 2.5% and 2.4% for DMG; 2.6% and 2.6% for methionine; and 3.1% and 3.4% for TMAO. Serum 5-methyltetrahydrolate (5-MeTHF) and folic acid were quantified in the Caudill laboratory using LC-MS/MS stable-isotope dilution methods [50] with modifications based on the instrumentation [51]. Intraassay and inter-assay CVs of in -house controls were 1.8% and 1.9% for 5-methyltetrahydrofolate; and 4.9 and 8.5% for folic acid.

### Differential gene expression

After normalization, surrogate variable analysis (SVA) was used to estimate and adjust for hidden cofounding variables on gene expression [52]. Differential gene expression was identified using the limma R package by a linear model that included the children’s diagnosis outcome and all surrogate variables [53]. Differential gene expression analysis with children diagnosed as ASD and children diagnosed as Non-TD were included in the same model with three levels of diagnosis [53]. Fold change and standard error were calculated using the limma R package. Differentially expressed transcripts were identified as those with an unadjusted *p*-value < 0.05. Genome-wide significant differentially expressed transcripts were classified as those with a false discovery rate (FDR) adjusted *p*-value (*q*-value) < 0.05.

### Gene ontology (GO) term and pathway enrichment analysis

Transcripts with significant expression levels or selected gene lists were exported to DAVID bioinformatics software with default settings for GO analysis [54,55]. The analysis was done using the GO ontology database and Fisher’s exact test with multiple test correction by the FDR method [55]. GO term enrichments were presented by hierarchical terms. GO terms with an FDR *q*-value < 0.05 were considered statistically significant.

### Weighted Gene Co-Expression Network Analysis (WGCNA)

A weighted gene co-expression network was built using the WGCNA R package [56,57] with normalized expression levels after adjustment for batch effects using the ComBat function from the sva R package [58]. The correlation matrix included all probes and all samples. To construct a signed adjacency matrix, estimated soft thresholding power was used with approximately scale-free topology (R^2 fit more than 0.8). Adjacency values were transformed into a signed topological overlap matrix (TOM). Co-expression modules were identified from a dissimilarity matrix (1-TOM) with a minimum module size of 30 probes. Module eigengenes were clustered on correlation. Similar modules were merged based on a cut height of 0.25 to generate co-expression modules. Each module’s expression profile was summarized into a module eigengene (ME) using the matched module’s first principle component. The correlation between each gene in the module with the ME was represented as intramodule connectivity (kME). Module hub probes were defined as the probe in each module with the highest module membership. Pearson’s correlation coefficient was used to measure the correlation between traits and modules.

### Cell type proportion deconvolution

CIBERSORT was used to estimate the proportions of each cell type using the default settings and the LM22 adult peripheral blood signature gene expression profiles [59]. Normalized expression levels adjusted for batch effects were used to estimate cell type proportions. Both relative and absolute modes were performed together with 100 permutation tests. *P*-values were calculated using FDR multiple test adjustment. Significant associations were defined based on FDR *q*-value < 0.05.

## Results

### Study sample characteristics and nutrient measurements

High quality RNA was isolated from 300 maternal peripheral blood samples collected during pregnancy in the MARBLES high risk ASD cohort (**Supplementary Table 1**). Children from MARBLES pregnancies were diagnosed at 3 years old as ASD (67, including 47 male and 20 female), Non-TD (79, including 46 male and 33 female), and TD (154, including 79 male and 75 female) (**Table 1**).

**Table 1.**
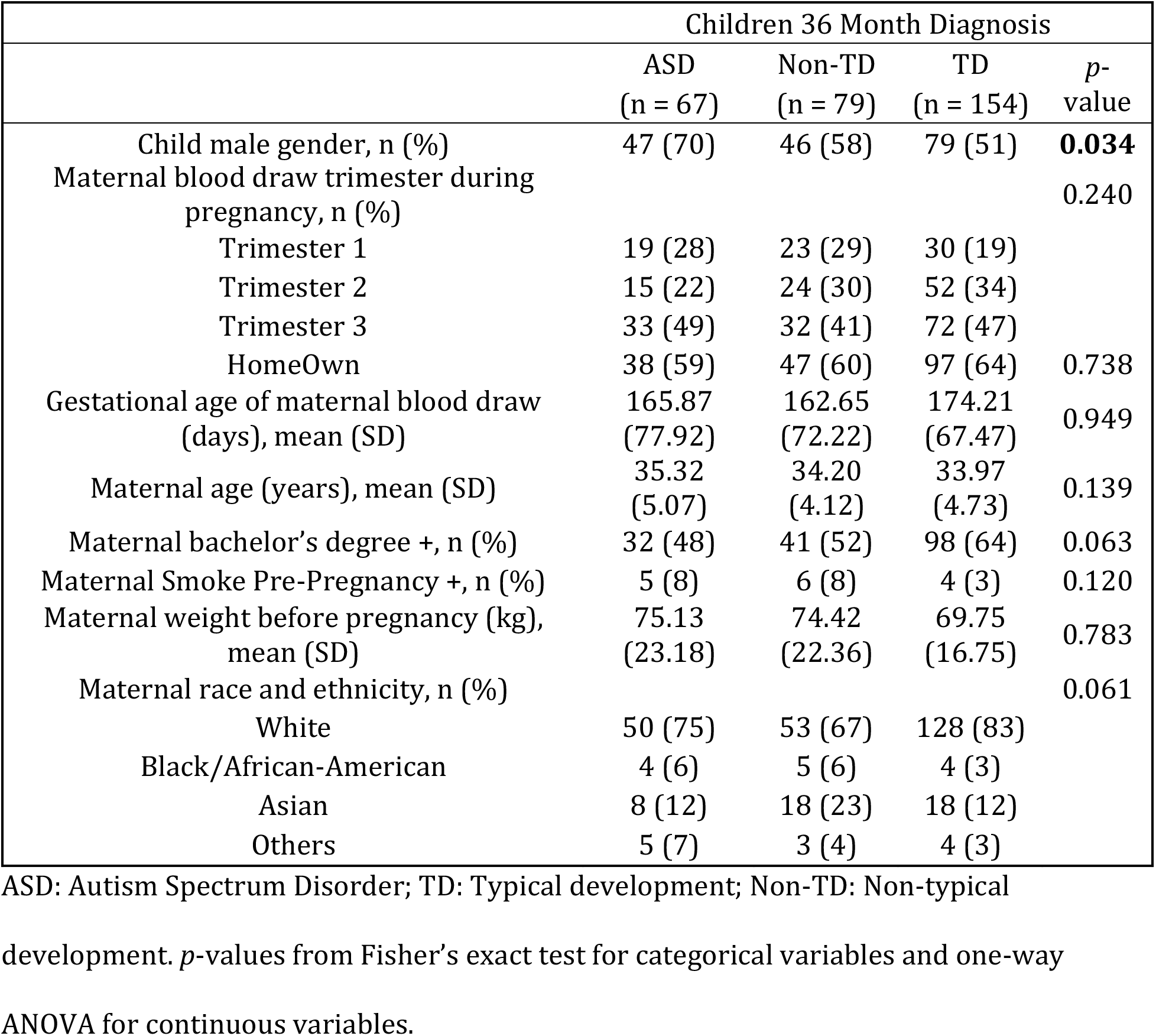
Demographic characteristics of mother participants and their children in MARBLES, stratified by child diagnosis outcomes.

Nutrients in the one-carbon metabolism pathway, including methionine, SAM, SAH, adenosine, homocysteine, 5-MeTHF, folic acid, Vit B6, Vit B12, choline, DMG, betaine, cystine, cysteine, GSH, and GSSG were directly measured from maternal blood in 14% – 62% of all collected samples (**Table 2**). None of these metabolites in maternal blood were significantly associated with clinical outcomes of children (**Table 2**).

**Table 2.**
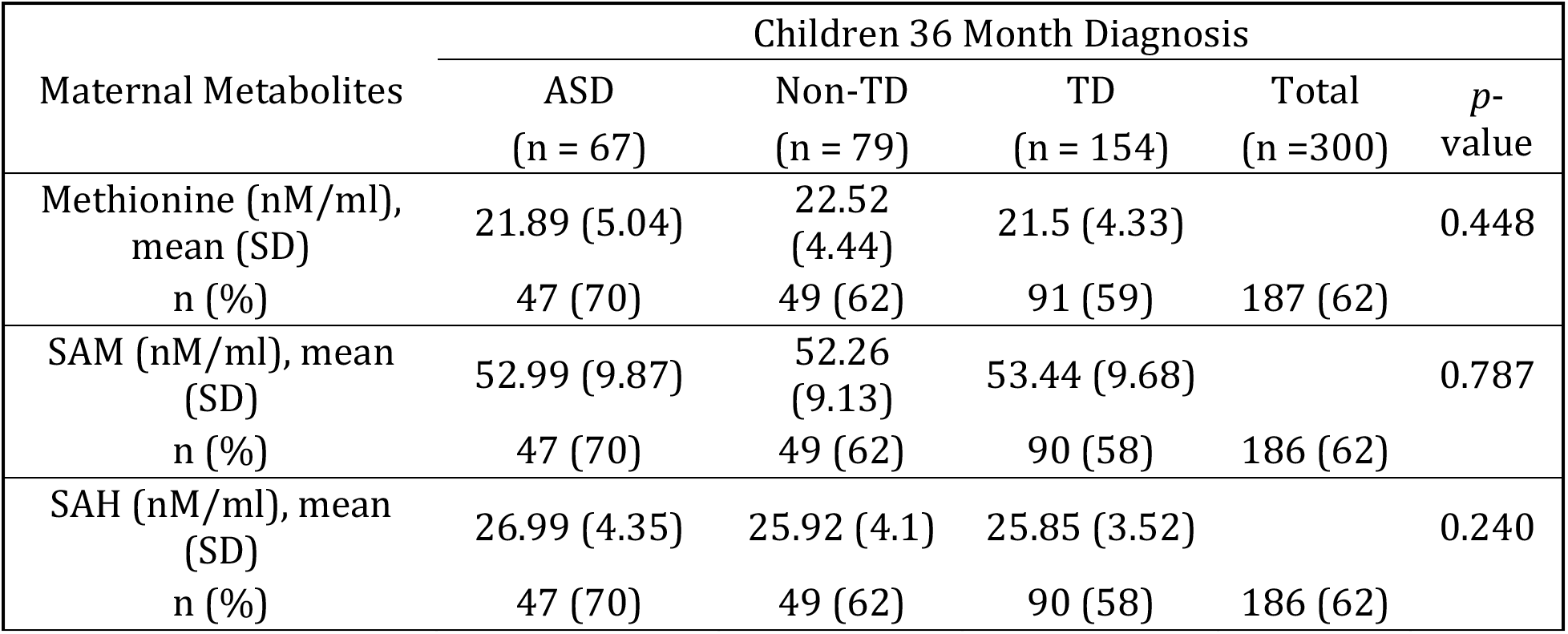
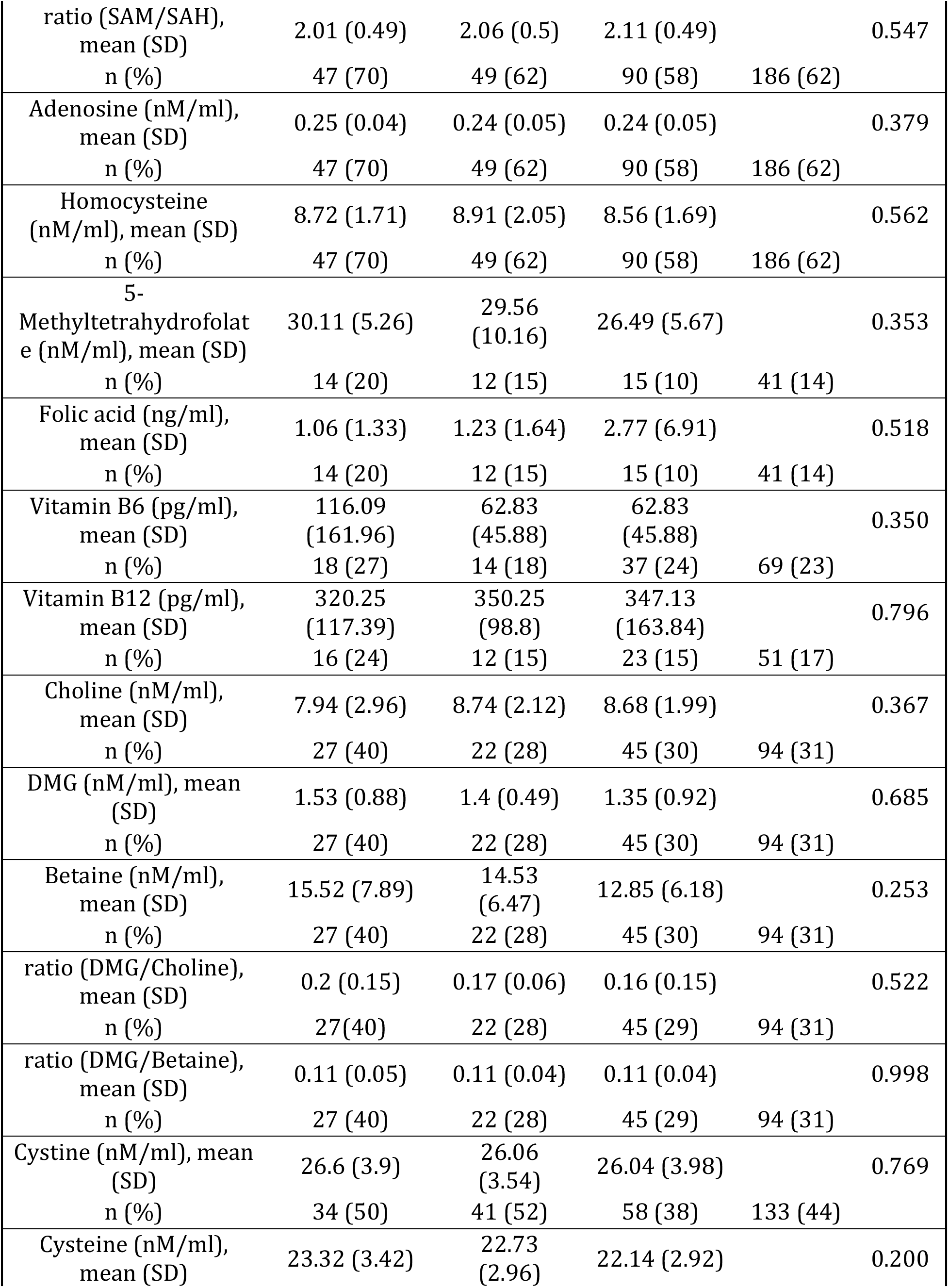
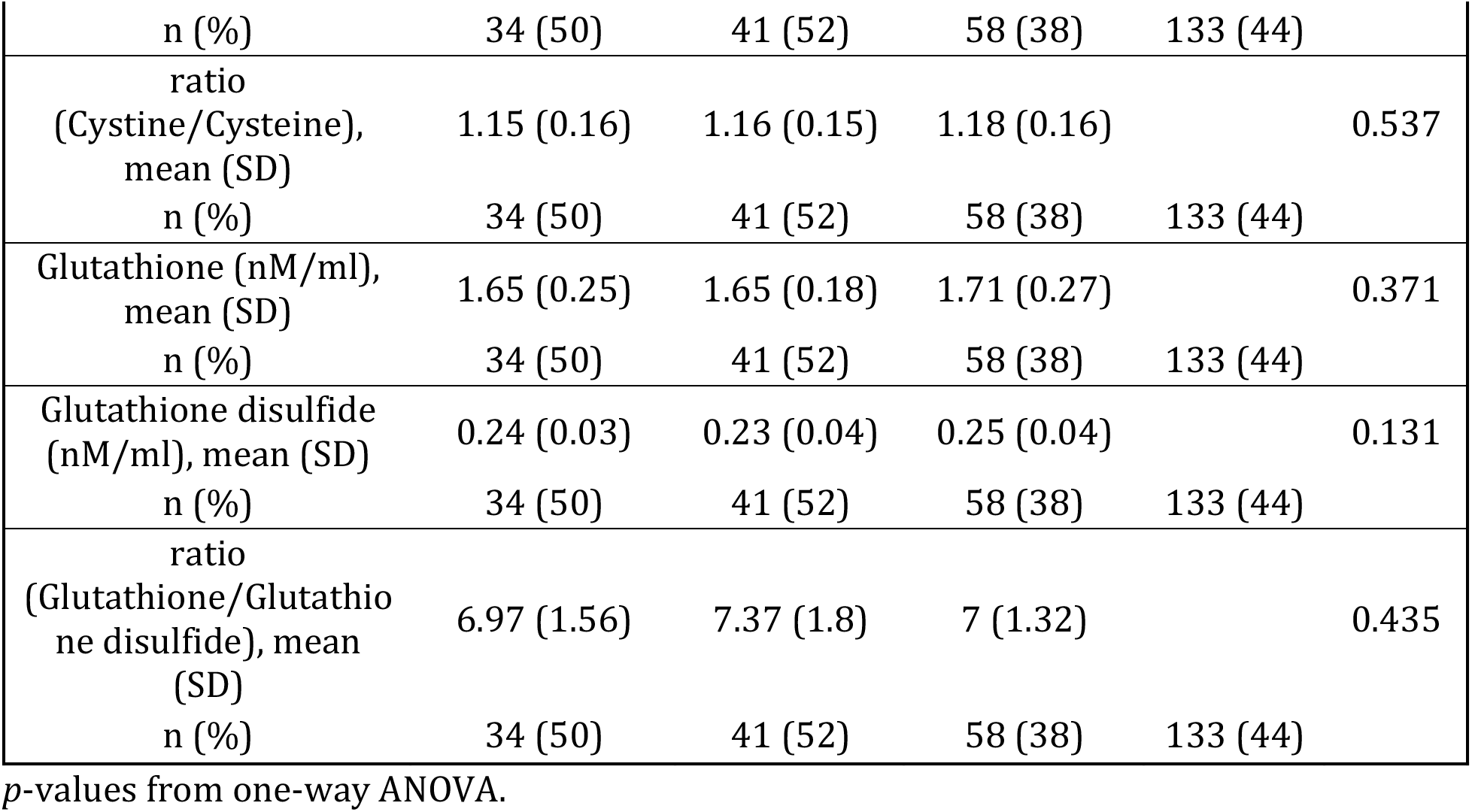
Descriptive statistics of maternal peripheral blood nutrients level in MARBLES, stratified by children diagnosis.

### Differential gene expression analyses by child outcome

Expression was measured using Human Gene 2.0 Affymetrix microarray and adjusted for surrogate variables, followed by differential gene expression analysis for child diagnosis (ASD, Non-TD, TD) on 36,459 transcripts. There were 28 surrogate variables (SVs) identified, including some significantly associated with batch effect and gestational age of maternal blood draw (**Supplementary Fig. 1**). Six transcripts located at four genes *(TGR-AS1, SQSTM1, HLA-C,* and *RFESD)* showed genome-wide significance with child outcomes (FDR adjusted *p*-value < 0.05) (**Supplementary Table 2**). Three out of six transcripts mapped to *HLA-C* (Major Histocompatibility Complex, Class I, C) (FDR adjusted *p*-value < 0.05). For those three *HLA-C* transcripts, increased levels were observed in ASD vs TD as well as Non-TD vs TD (unadjusted *p*-value < 0.05) (**Supplementary Fig. 2**).

Comparing the maternal blood transcriptome between ASD and TD outcomes revealed 2,012 differentially expressed transcripts at a marginal confidence level (unadjusted *p*-value < 0.05) that mapped to 1,912 genes, including 980 up-regulated and 1,032 down-regulated transcripts, with none significant after FDR adjustment (**Fig. 1A**, **Supplementary Table 3**). There was a significant overlap between these 1,912 differentially expressed genes and a list of strong ASD candidate genes from the Simons Foundation Autism Research Initiative (SFARI Gene, including *TRIO, GRIA1, SMARCC2, SPAST, DIP2C, FOXP1, CNTN4,* Fisher’s exact test, *p*-value < 0.05) [60].

**Figure 1.**
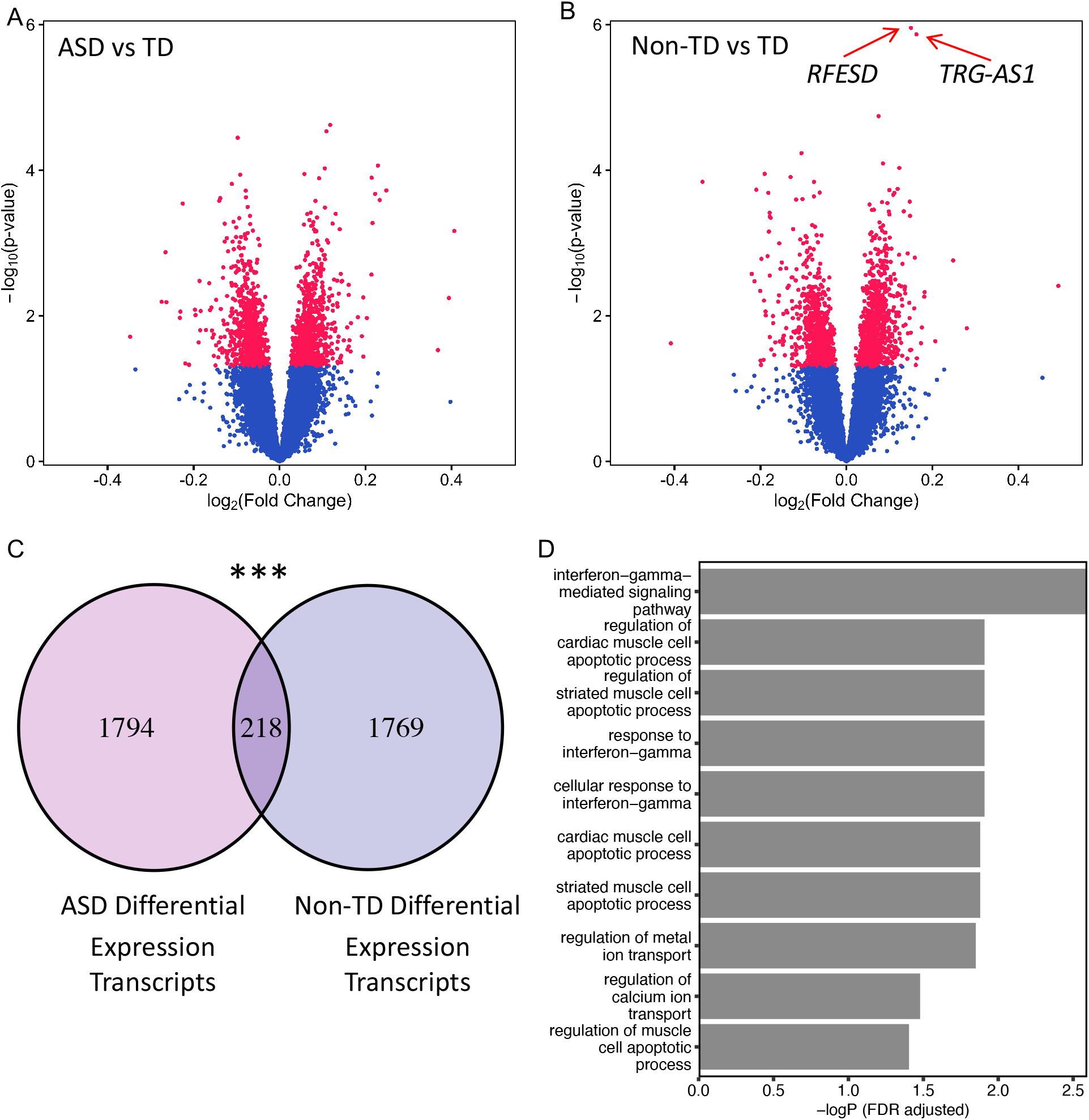
Identification and function of ASD associated and Non-TD associated differentially expressed genes in maternal peripheral blood. Differential expression analysis was performed in maternal peripheral blood transcriptomes (n = 300) after adjustment for surrogate variables. A) Identification of 1,912 differentially expressed genes (2,012 transcripts, *p*-value < 0.05) compared between children diagnosed as ASD (n = 67) and TD (n = 154). B) Identification of 1,919 differential expressed genes (1,987 transcripts, *p*-value < 0.05) compared between children diagnosed as Non-TD (n = 79) and TD (n =154). Two transcripts located at *RFESD* and *TRG-AS1* were genome-wide significant in the Non-TD to TD comparison and at an unadjusted *p* value in ASD to TD comparison (Supplementary Table 2). C) Venn diagram represents the overlap in differentially expressed transcripts (unadjusted p<0.05) identified in ASD to TD versus Non-TD to TD comparisons, which was greater than expected by random using a Fisher’s exact test (*p*-value < 0.001***). D) Gene ontology (GO) and pathway analysis was performed on the 218 transcripts differentially expressed in both ASD-TD and non-TD-TD comparisons, with significant enrichments (Fisher’s exact test, FDR *p*-value < 0.05). In contrast, the differentially expressed transcripts uniquely associating with either ASD or non-TD were not significantly enriched for any GO terms.

Comparing the maternal blood transcriptome between Non-TD and TD outcomes revealed 1,987 differentially expressed transcripts at a marginal confidence level (unadjusted *p*-value < 0.05) that mapped to 1,919 genes, including 1,044 up-regulated and 943 down-regulated transcripts (**Fig. 1B**, **Supplementary Table 4**). Two of these transcripts at *RFESD* and *TRG-AS1* genes also passed genome-wide significance (FDR adjusted *p*-value < 0.05). Unlike the ASD vs TD comparison, however, no significant overlap was observed between Non-TD differentially expressed genes and SFARI gene lists.

A significant overlap of 218 transcripts was observed between ASD associated differentially expressed transcripts and Non-TD associated differentially expressed transcripts (Fisher’s exact test, *p*-value < 2.2E-16) (**Fig. 1C**). Gene ontology (GO) analysis of these 218 transcripts revealed significant enrichment for the interferon-gamma mediated signaling pathway, apoptosis in muscle, response to interferon gamma, and metal ion transport (**Fig. 1D**, **Supplementary Fig. 3**). CaMK (calmodulin-dependent protein kinase)families *(CAMK2A, CAMK2B, CAMK2D* and *CAMK2G)* and HLA (human leukocyte antigen) systems *(HLA-B, HLA-C* and *HLA-E)* were included in those pathways (**Fig. 1D**). In contrast, neither list of ASD-or TD-specific differentially expressed transcripts were significantly enriched for any GO terms.

### Weighted gene co-expression network analysis (WGCNA) identified gene modules correlating with specific maternal nutrient levels

WGCNA was performed as a complementary bioinformatic approach that incorporates the independent and inter-related associations of transcript levels with measured concentrations of maternal nutrients. First, expression values were adjusted for batch effect, then correlation patterns among all transcripts were analyzed across all 300 samples. WGCNA identified 27 co-expressed gene modules in our dataset, representing 17,049 transcripts, distinguished from 19,410 transcripts not clustered and grouped into the “grey” module (**Fig. 2A, Fig. 2B**, **Supplementary Fig. 4**, **Supplementary Table 5**). For each module, the number of transcripts, as well as the hub gene, defined as the gene with the highest correlation with the module eigengene, were determined (**Fig. 2B**, **Supplementary Table 6**). Out of those 27 co-expression modules, 23 modules showed associations between eigengene expression level and at least one variable related to demographics, diagnosis, or maternal nutrients, after correction (FDR adjusted *p*-value < 0.05) (**Fig. 2A**, **Supplementary Fig. 4**). All 27 modules were significantly associated with one or more covariates at unadjusted *p*-value < 0.05 (**Fig. 2A**, **Supplementary Fig. 5**).

**Figure 2.**
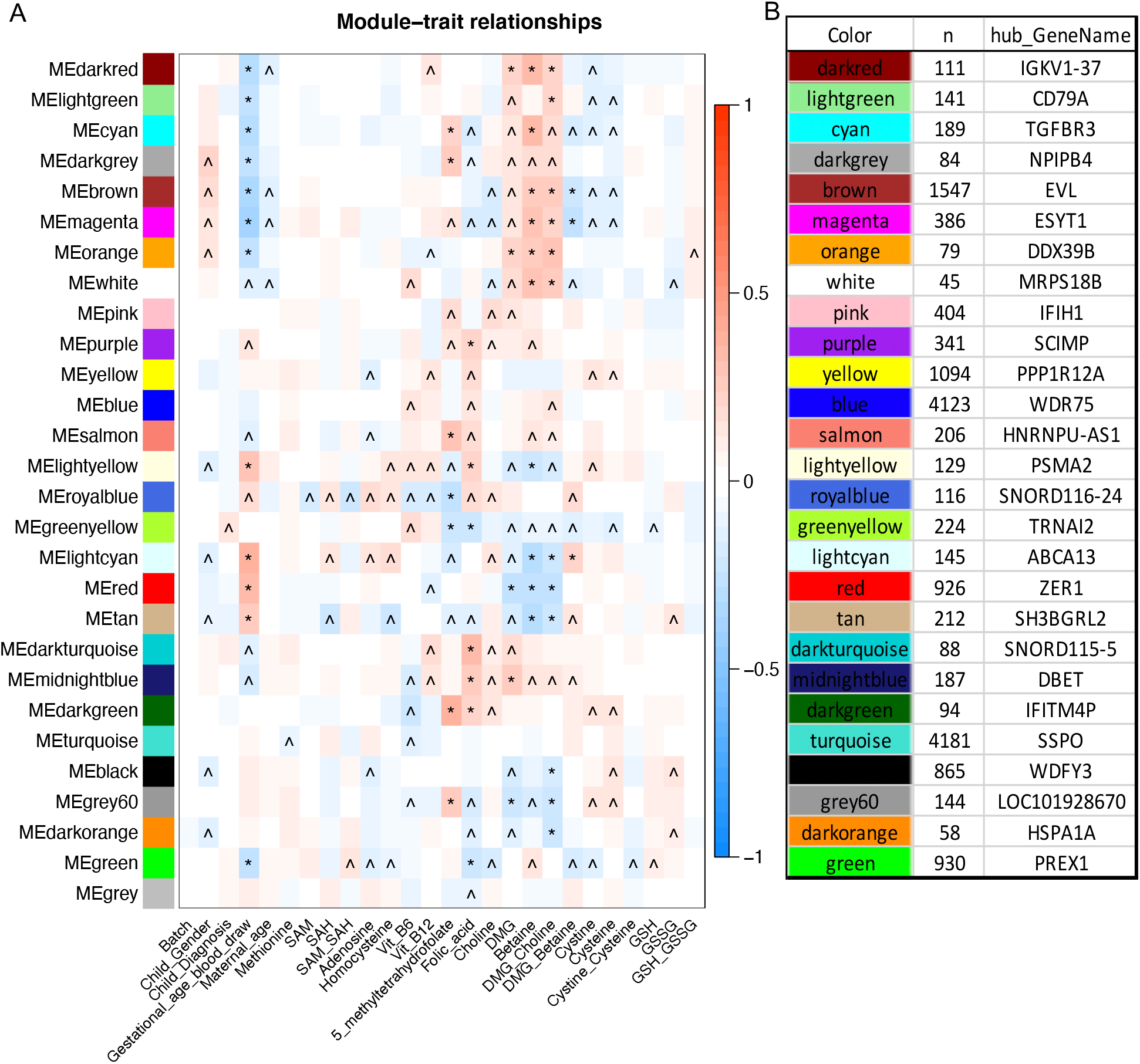
Co-expression network modules with demographic factors and maternal peripheral blood one-carbon metabolites. A) Heatmap of Z-scores of modules eigengenes with sample covariates with 27 coexpression network modules on all 300 maternal blood samples. Each row represents a different module eigengene and each column is the associated trait, which include child clinical outcome, demographic factors, and maternal blood metabolite concentrations. The matrix was calculated by Pearson correlation and *p-*values adjusted for the total number of comparisons. Color represents the direction (red, positive correlation; blue, negative correlation) and intensity reflects the significance. (^ unadjusted *p*-value < 0.05 and FDR adjusted *p*-value > 0.05; * FDR adjusted *p*-value < 0.05) B) Number of transcripts and hub genes from all 27 co-expressed modules are listed.

Multiple co-expression modules were significantly correlated (FDR adjusted *p*-value < 0.05) with gestational age at blood draw and four maternal metabolites, including 5-MeTHF, folic acid, DMG, and betaine (**Fig. 2A**). None of the additional measured variables was associated with any co-expression gene modules at a high confidence level, including clinical outcome. However, the module “greenyellow” showed a marginally significant positive correlation with outcome (unadjusted *p*-value = 0.02, FDR adjusted *p*-value = 0.14) and a negative significant correlation with both 5-MeTHF (FDR adjusted *p*-value = 0.02) and folic acid levels (FDR adjusted *p*-value = 0.02) (**Fig. 2A, Fig. 3A, Fig. 3B, Supplementary Fig. 4, Supplementary Fig. 5**, **Supplementary Table 7**). This greenyellow module eigengene also showed opposite correlation between ASD and 5-MeTHF, consistent with 5-MeTHF protective functions in ASD (**Fig. 3A**, **Fig. 3B**).

**Figure 3.**
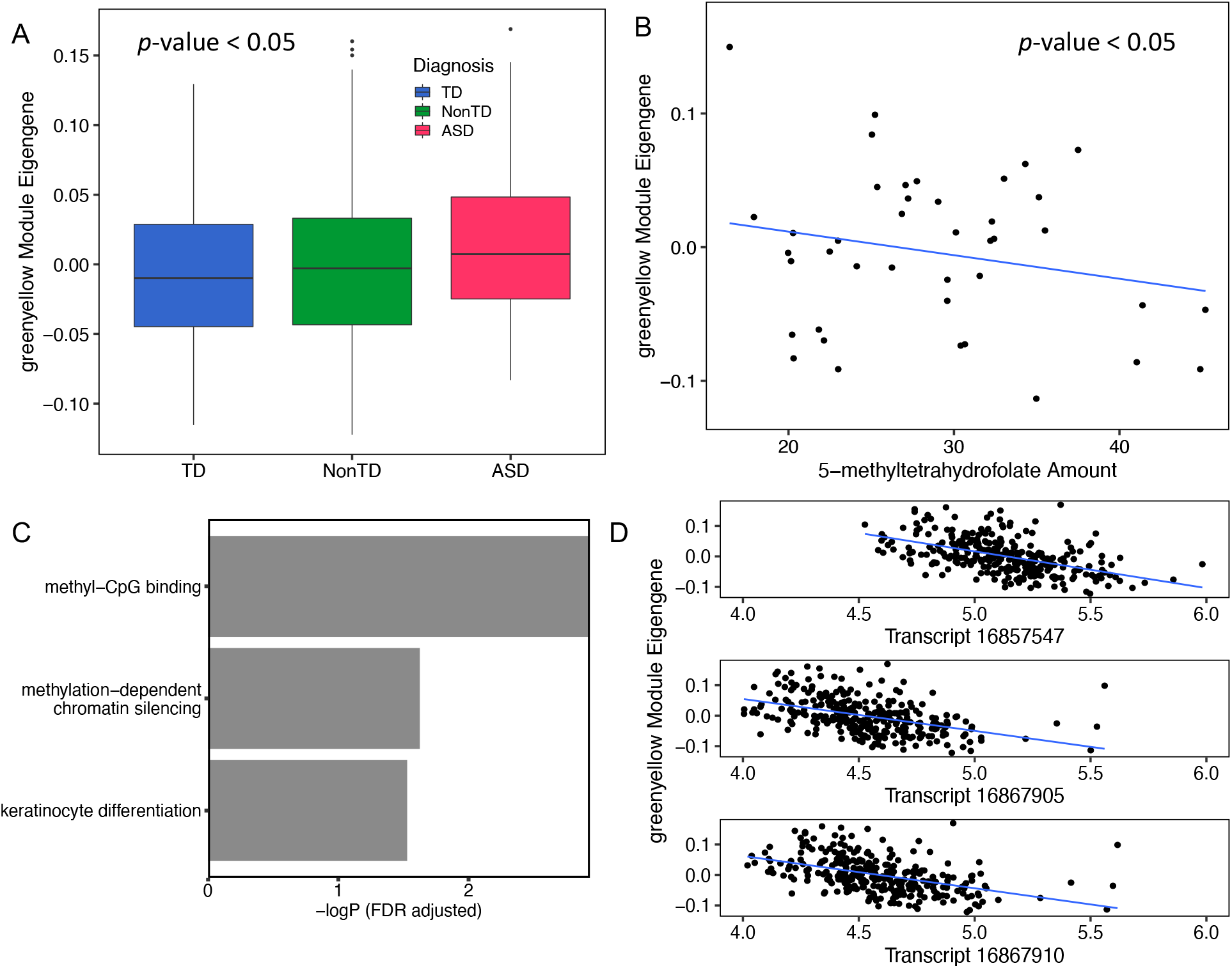
“Greenyellow” module was positively associated with diagnosis and negatively associated with folic acid and 5-MeTHF. A) “Greenyellow” module eigengene was significantly associated with child diagnosis (one way ANOVA, unadjusted *p*-value < 0.05). Greenyellow eigengene values were higher in maternal blood from ASD pregnancies than TD or non-TD pregnancies. B) “Greenyellow” module eigengene level was significantly negatively associated with 5-MeTHF concentrations in maternal blood (ANOVA, *p*-value < 0.05). C) Bar graph shows gene ontology (GO) and pathway significant enrichments from the 224 transcripts in “greenyellow” module (Supplementary Table 8). D) Transcripts (16857547, 16867905 and 16867910) from MBD3L3-5 genes encoding proteins involved in methylation-CpG binding functions were significantly negatively associated with “greenyellow” module eigengene.

This “greenyellow” module contained 224 transcripts with *TRNAI2* as hub gene (**Supplementary Table 7**). These 224 transcripts showed a significant enrichment for gene ontology functions in methylation-CpG binding, methyl-dependent chromatin silencing, and keratinocyte differentiation (Fisher’s exact test, FDR adjusted *p*-value < 0.05) (**Fig. 3C**, **Supplementary Table 8**). The three known genes with methyl-binding functions included *MBD3L3, MBD3L4* and *MBD3L5,* represented by 16857547, 16867905, and 16867910 transcripts (**Fig. 3C**). Normalized expression of those three transcripts was also significantly associated with the “greenyellow” module eigengene, supporting their membership in the module (**Fig. 3D**).

### Eight co-expression modules strongly clustered with betaine and DMG

Among the 27 identified co-expression modules, eight modules (darkred, lightgreen, cyan, darkgrey, brown, magenta, orange and white) were highly correlated with each other and clustered based on unsupervised hierarchical clustering, representing a total of 2,582 transcripts (**Fig. 4, Supplementary Fig. 6, Supplementary Table 9**). Betaine and DMG were significantly associated and clustered together with this distinct block of coexpression modules.

**Figure 4.**
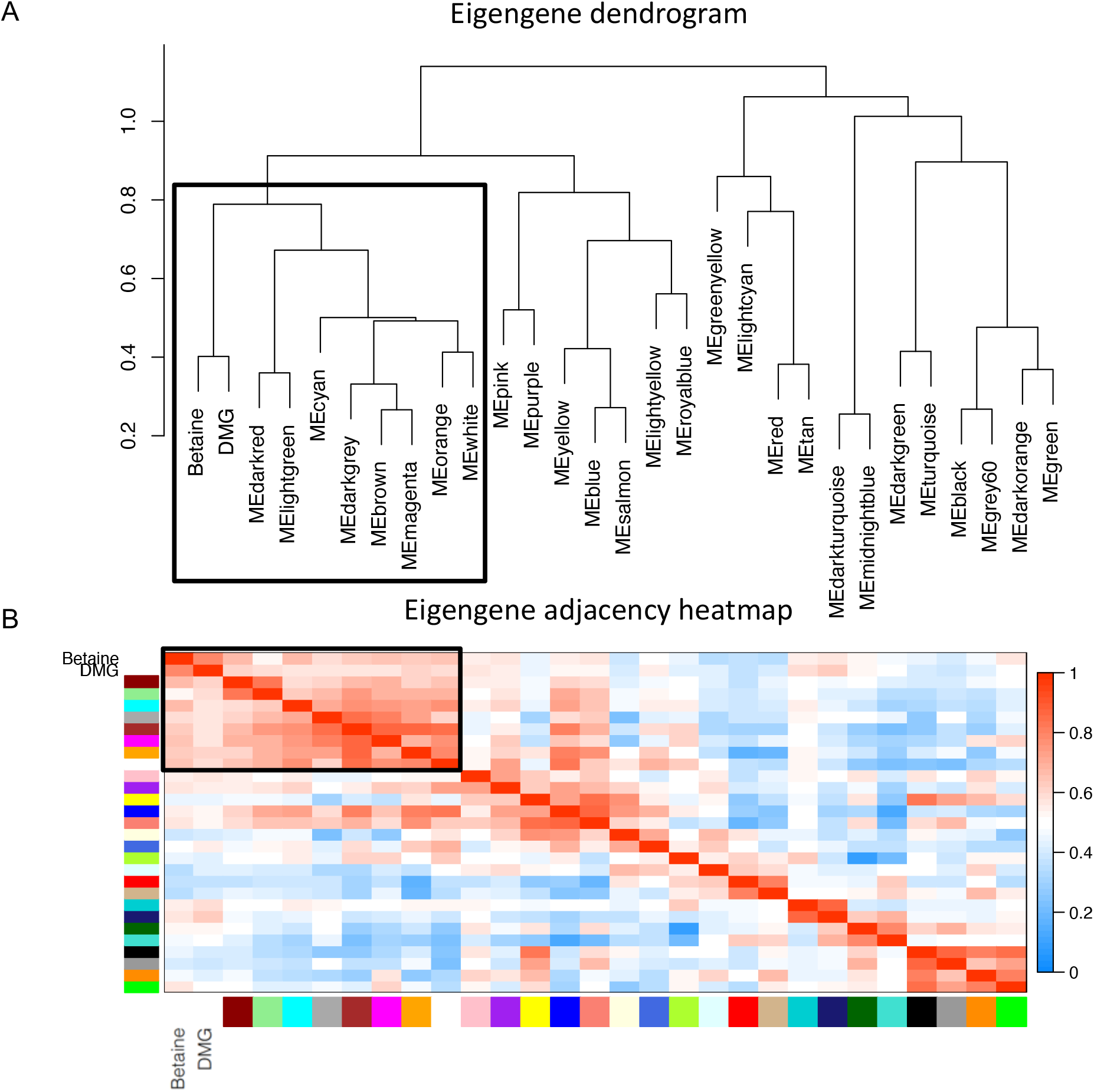
Eight weighted gene co-expression modules associated with maternal betaine and DMG concentrations were strongly clustered. A) Unsupervised hierarchical clustering dendrogram was performed with module eigengenes, betaine and DMG. The height of each node represents the intergroup dissimilarity. Similar nodes clustered together under one branch. B) Unsupervised hierarchical clustering adjacency heatmap, with color and intensity representing the degree of correlation (dark, high; light, low correlation). Black box indicates the block of eight weighted gene co-expression modules associated with betaine and DMG concentrations.

Transcripts inside these eight clustered co-expression modules associated with betaine and DMG showed significant enrichment for 18 gene pathways involved in adaptive immune response, RNA processing, histone modification, inflammatory response, and Rett syndrome (Fisher’s exact test, FDR adjusted *p*-value < 0.05) (**Supplementary Fig. 7**). Network analysis using GeneMANIA [61] identified a network with *EVL* in the center, linked with other hub genes (**Supplementary Fig. 8**).

### Cell type composition in maternal peripheral blood associated with maternal metabolites but not child clinical outcomes

In order to determine the effects of cell composition differences on the findings associated with maternal transcriptomes, cell type specific information from 22 immune cell types was deconvoluted using the CIBERSORT web tool. Maternal peripheral blood samples reflected a mixture of cell types, with neutrophils as the largest and most variable population ranging from 17% to 48% (**Fig. 5A, Supplementary Table 10**). The eigengenes for 21 out of 27 modules were significantly correlated with at least one cell type (FDR adjusted *p*-value < 0.05) (**Supplementary Fig. 9**). No significant difference was observed in cell type composition between child diagnosis outcomes or gender (**Fig. 5A. 5B**, **Supplementary Fig. 10**). Furthermore, neither the “greenyellow” module, nor the betaine and DMG variables were significantly associated with cell type proportions, suggesting that the associations identified with these modules were largely cell type independent (**Fig. 5B**, **Supplementary Fig. 9**, **Supplementary Fig. 10**).

**Figure 5.**
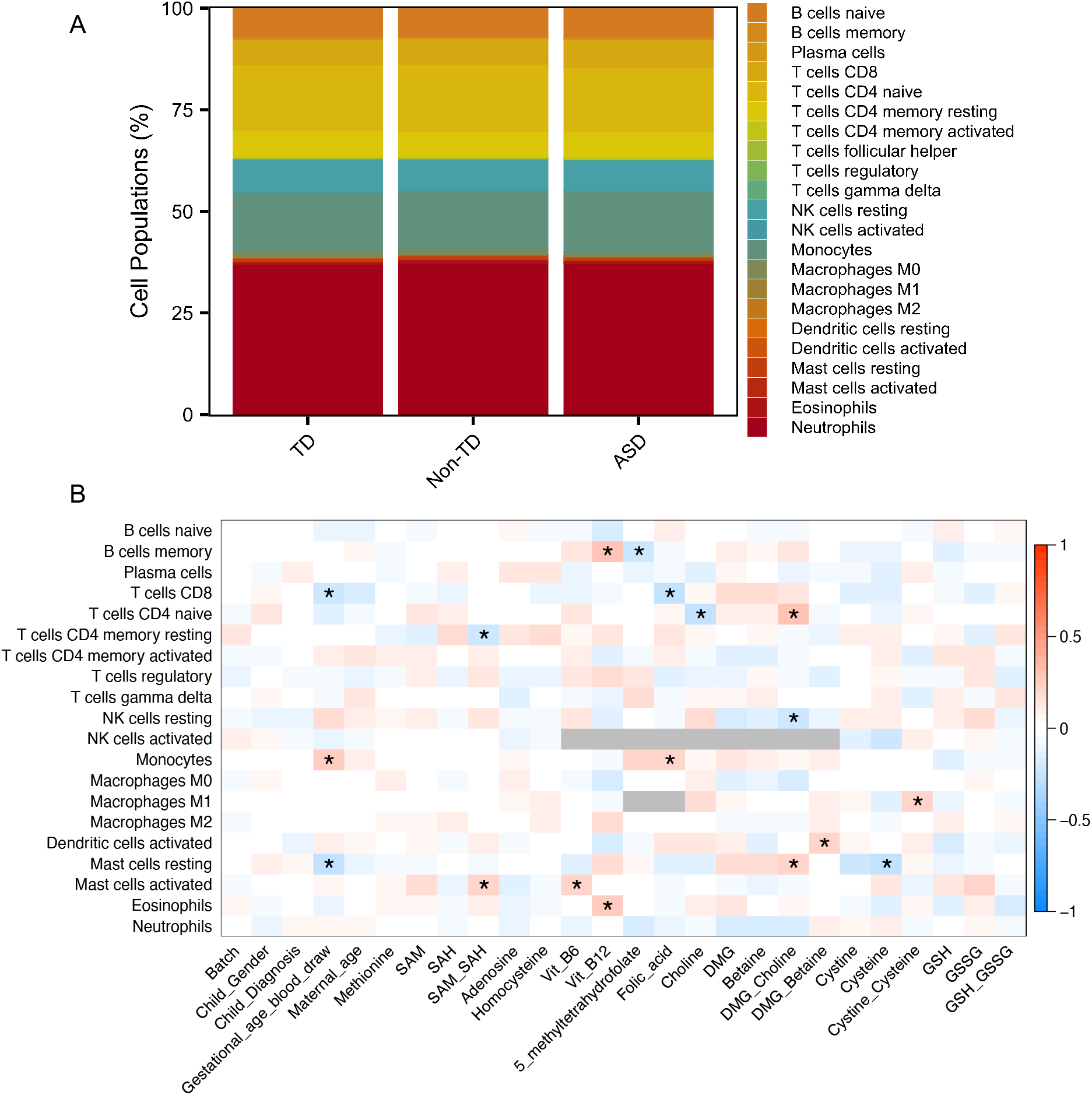
Imputed cell type proportions in material peripheral blood associated with demographic factors and maternal nutrients. A) Barplot of each cell type mean estimated proportion separated by children diagnosis outcomes using peripheral blood reference panel in CIBERSORT. B) Heatmap of correlation between sample demographic factors and maternal nutrients with cell type proportions. Each row represents a cell type proportion and columns represent traits, including child diagnostic outcome, demographic factors, and maternal blood nutrient concentrations. *p-*values adjusted for the total number of comparisons. Color represents the direction (red, positive correlation; blue, negative correlation) and intensity reflects the significance, * *p*-value < 0.05 after FDR correction.

In contrast, some cell type proportions were significantly correlated with some maternal metabolites. Vit B6, 5-MeTHF, choline, cysteine, the ratio of DMG/betaine, and the ratio of cystine/cysteine were separately associated with six cell types (FDR adjusted *p*-value < 0.05) (**Fig. 5B, Supplementary Fig. 10**). Vit B12, folic acid, the ratio of DMG/betaine, and the ratio of SAM/SAH were associated with more than one cell type (FDR adjusted *p*-value < 0.05) (**Fig. 5B, Supplementary Fig. 10**). The most significant association was between vit B12 and memory B cells (FDR adjusted *p*-value = 0.0001) (**Fig. 5B, Supplementary Fig. 10**).

## Discussion

Maternal blood collected during pregnancy provides molecular insights into the *in utero* environment relevant to the etiology of ASD. This was the first study to our knowledge to examine gene expression differences in maternal peripheral blood during pregnancy together with *in utero* maternal one-carbon metabolites for children that went on to develop ASD, TD, or Non-TD at 36 months. Using complementary bioinformatics approaches, we identify several genes and gene pathways consistent with proinflammatory and oxidative stress responses in mothers of children with adverse neurodevelopmental outcomes. We also identify eight novel coregulated gene modules associated with maternal blood betaine and DMG concentration.

### Genes and gene patterns common to mothers of children with ASD and non-typical neurodevelopment

Using differential gene expression analysis of individual genes, we describe four genes *(SQSTM1, HLA-C, TRG-AS1, RFESD)* that were differentially expressed in mothers of TD children compared with those diagnosed as either ASD or non-TD. *SQSTM1* encodes the p62 sequestosome that acts as a receptor for ubiquitinated cargo in the selective autophagy response induced by oxidative stress [62], and also links mTOR and GABA signaling pathways in brain [63]. *RFESD,* encoding an iron-sulfur cluster binding protein with oxidoreductase activity, is located on 5q15, a hotspot for copy number variants in intellectual and developmental disabilities [64,65]. *TRG-AS1,* T-cell receptor gamma locus antisense RNA 1, is located on 7p14.1, another locus previously associated with developmental delay, intellectual disability, and ASD [64,66,67]. *HLA-C* belongs to the HLA (human leukocyte antigen) polymorphic loci encoding major histocompatibility class I (MHC I) proteins involved in antigen presentation to CD8+ T cells and NK cells. HLA-C is important for both tolerance to fetal allo-antigens and viral immunity during pregnancy [68]. Proinflammatory cytokines such as interferon gamma (IFNγ) induce HLA-C expression in both lymphocytes and placental trophoblasts. A number of previous studies have shown that the HLA locus is associated with ASD [69–71] or HLA locus activation in ASD children and their mothers[72–74], which is consistent with our findings of elevated *HLA-C* expression levels in ASD and Non-TD compared to TD mothers. Furthermore, two additional class I loci, *HLA-B* and *HLA-E,* were also differentially expressed in mothers of children classified as ASD and Non-TD compared to TD children in this study, providing further evidence of an MHC I response in pregnancies of atypical neurodevelopment.

Furthermore, gene pathway analysis of differentially expressed genes common to nontypical development revealed enrichment for the interferon-gamma mediated signaling pathway, which has been previously found to be elevated in mothers of children with ASD and other neurodevelopmental disorders [75,76]. In one such study, elevated interferongamma levels in maternal midgestation peripheral blood was associated with a 50% increased risk of offspring ASD risk [75]. A second enriched pathway included CaMK family members which play an important role in neuronal connectivity and synaptic plasticity [10,77,78] as well as immune response and inflammation [79]. Prior ASD studies have implicated the CaMK pathway in the dendritic growth and local connectivity alterations related to gene-environment interactions in ASD [10,77,78].

While genome-wide significance of individual differentially expressed genes was not observed between samples from mothers whose children developed ASD compared to TD after adjusting for multiple comparisons, seven genes with significant unadjusted *p* values were also on the SFARI list of strong ASD candidate genes. *TRIO,* Trio Rho guanine nucleotide exchange factor, promotes exchange of GDP for GTP and provides necessary support for cell migration and cell growth related to Alzheimer disease and other types of neurological conditions [80,81]. *GRIA1,* the predominant excitatory neurotransmitter in brain, is associated with the activation of normal neurophysiologic processes [80,82].

*SMARCC2* encodes a chromatin remodeling protein with helicase and ATPase activities which has been implicated in altering chromatin structure in ASD [83]. *CNTN4* functions in neuronal network formation and plasticity, and is associated with nervous system development at the transcriptome level [84,85]. Mutations in *FOXP1,* a developmental transcription factor, are observed in rare cases of intellectual disability with ASD [86,87].

### Methylation and methyl-binding functions in a gene module oppositely associated with folic acid and ASD risk

The complementary co-expression network analysis further revealed a module of 224 coexpressed genes in maternal blood showing an association with folic acid and 5-MeTHF levels in the opposite direction from ASD risk that could not be explained by cell type differences. Interestingly, these ASD and nutrient associated genes were functionally enriched for DNA methylation binding and methylation-dependent chromatin silencing, consistent with prior DNA methylation changes observed in ASD [88–91] as well as ASD-like syndromes associated with methyl binding proteins [92,93]. Folic acid, the synthetic form of folate that contributes substrate for one-carbon metabolism, and 5-MeTHF, one of the active biological forms of folate that plays a critical role in one-carbon metabolism, have also been shown to be inversely associated with developmental delay [14,15,94,95]. *MBD3L* is predicted to assist with demethylation reactions and functions as a transcriptional repressor [96–98].

One-carbon metabolites associated with changes in gene expression in this study have also been associated with prevention of numerous conditions [89,99–101]. The coregulated block of betaine and DMG co-expression modules contained genes enriched in the adaptive immune system and chromatin modification functions, as well as Rett syndrome, a known syndromic form of ASD [102–105]. Choline can be metabolized to betaine, which converts homocysteine to form methionine and generates DMG in the one-carbon pathway [106,107]. A previous study of maternal peripheral blood collected at term showed that changes of betaine and DMG were in the opposite direction from choline when compared with nonpregnant women [108]. *EVL* (Enah/Vasp-like) is involved in actin cytoskeleton remodeling and is crucial for central neuron system processes and immune system functions [109–111]. One study also showed *EVL* as a differentially expressed gene in schizophrenia in peripheral blood [110].

Previous studies in ASD have been focused on post mortem brain tissue [28,112], as a tissue relevant to the disorder, but collected after diagnoses were made, raising concerns about reverse causation in determining etiologically-relevant expression changes. Few studies have focused on prospective transcriptomic profiles collected prior to the presentation of the disorder [31,113], and none have examined maternal gene expression profiles during pregnancies at high risk for developing ASD. In addition, few studies have integrated maternal transcriptome and one-carbon metabolite data within biospecimens. Furthermore, most studies of ASD expression biomarkers have not considered the roles of nutritional factors during pregnancy that could be relevant to fetal development.

## Limitations

A limitation of using maternal peripheral blood to examine expression is that it contains multiple cell types, and proportions can differ across samples. However, estimated cell type composition of maternal blood was not significantly associated with the child’s clinical outcomes, the “greenyellow” module, betaine, or DMG, which suggests that our main findings were not driven by differential cell type proportions. After correcting for multiple comparisons, this study did not identify any individual differentially maternally expressed genes specifically associated with ASD, although 6 transcripts from 4 genes reached genome-wide significance with diagnosis of either ASD or Non-TD. Furthermore, lack of genome-wide evidence of individual differentially expressed genes specific to a pairwise comparison of ASD vs. TD is likely due to the relatively small sample size that is inherent to a prospective ASD study, but likely underpowered to detect small differences in transcript levels. However, this does not eliminate the importance of identifying and understanding the biologically significant gene set enrichments and co-expression network modules using differential expression gene and WGCNA analysis. Additionally, other factors, including genetics, epigenetics, and other environmental factors can influence the transcriptome and ASD risk. Approaches incorporating those factors will be important in future studies.

## Conclusions

In summary, genome-wide gene expression analysis of maternal peripheral blood samples revealed transcriptome changes associated with maternal one-carbon metabolites and child neurodevelopmental outcomes implicating maternal immune, apoptotic, and epigenetic mechanisms in ASD. In addition, folic acid and 5-MeTHF were associated with expression of genes involved in methylated-CpG binding in an opposite direction to that of ASD, consistent with prior evidence of protection. Finally, maternal betaine and DMG levels clustered with co-expressed genes related to immune, chromatin modification, and development functions. These results therefore provide important biological insights into maternal gene pathways associated with adverse neurodevelopment in the child, as well as the protective role of one carbon metabolites in the complex etiology of ASD.

## List of abbreviations

ASD: autism spectrum disorder,
Non-TD: non-typical development,
TD: typical development,
ADOS: autism diagnostic observation schedule,
ADI-R: autism diagnostic interview – revised,
MARBLES: Markers of Autism Risk in Babies – Learning Early Signs,
WGCNA: weighted gene correlation network analysis,
FDR: false discovery rate,
SD: standard deviations,
SVA: surrogate variable analysis,
GO: gene ontology,
SFARI: Simons Foundation Autism Research Initiate,
5-MeTHF: 5’-methyltretrahydrofolate,
DMG: dimethylglycine,
SAM: S-adenosylmethionine,
SAH: S-adenosylhomocysteine,
GSH: glutathione,
GSSG: glutathione disulfide,
Vit B6: vitamin B6,
Vit B12: vitamin B12

## Declarations

### Ethics approval and consent to participate

The UC Davis Institutional Review Board and the State of California Committee for the Protection of Human Subjects approved this study and the MARBLES Study protocols. All data and specimens were collected after parent given written informed consent form.

### Consent for publication

Not applicable.

### Availability of data and material

Data are shared in the Gene Expression Omnibus (GEO) accession number (GSE148450) based on participates consent. Code and scripts for this study are available on GitHub (https://github.com/Yihui-Zhu/AutismMaternalBloodExpression). Other related data and information are included at supplementary tables.

### Authors’ contributions

YZ was the lead author and contributed substantially to all bioinformatics data analysis, data visualization, interpretation of results, and writing the manuscript. CEM and BPD added critical advice for bioinformatics data analysis methods and interpretation. MAC, OVM, JWM, JSJ and SBM contributed to nutrient metabolite measurements. MDF, IH and RJS contributed to study design, and subject acquisition, diagnosis and characterization. RJS and JML conceived the study and contributed substantially to data interpretation and revision of the manuscript. All authors read and approved the final manuscript.

## Acknowledgements

We would like to thank the UCD Children’s Center for Environmental Health for helpful discussions and the MARBLES study participants. We would also like to thank Daniel Young provided substantial assistance with multiple data sets data preparation.

## Additional Files

### Supplementary Figures

**Supplementary Figure 1.** Surrogate variable analysis in MARBLES subjects.

**Supplementary Figure 2.** Three transcripts at *HLA-C* reached genome-wide significance with diagnosis.

A, B and C show the expression level at each transcript. The y-axis shown the normalized and adjusted expression level. The x-axis represented three diagnosis groups, ASD, Non-TD and TD.

A) Transcript 17039281, F-test showed the significant association between expression and diagnosis (unadjusted *p*-value = 3.16E-06, FDR adjusted *p*-value = 0.038). For ASD compared with TD group, expression was significantly associated with ASD prior to genome-wide correction (unadjusted *p*-value = 5.17E-04, FDR adjusted *p*-value = 0.57). In Non-TD compared to TD, expression was also significant associated with Non-TD prior to genome-wide correction (unadjusted *p*-value = 0.017, FDR adjusted *p*-value = 0.81).

B) Transcript 17041782, diagnosis (unadjusted *p*-value = 6.07E-06, FDR adjusted *p*-value = 0.046); ASD vs TD (unadjusted *p*-value = 5.83E-04, FDR adjusted *p*-value = 0.57); Non-TD vs TD (unadjusted *p*-value = 0.025, FDR adjusted *p*-value = 0.84).

C) Transcript 17031781, diagnosis (unadjusted *p*-value = 7.96E-06, FDR adjusted *p*-value = 0.048); ASD vs TD (unadjusted *p*-value = 6.5E-04, FDR adjusted *p*-value = 0.57); Non-TD vs TD (unadjusted *p*-value = 0.027, FDR adjusted *p*-value = 0.86).

**Supplementary Figure 3.** Gene ontology and pathway analysis using directed acyclic graph (DAG) on the 218 transcripts common to ASD vs. TD and Non-TD vs. TD differentially expressed gene lists.

**Supplementary Figure 4.** Co-expression network modules with diagnosis, demographic factors, and maternal blood nutrient concentrations. The values in the cells represent Pearson r (adjusted *p*-value). *p*-value were adjusted for all comparisons.

**Supplementary Figure 5.** Co-expression network modules with diagnosis, demographic factors, and maternal blood nutrient concentrations. The values in the cells represent Pearson r (*p*-value). *p*-value shown here were unadjusted *p*-value without adjustment.

**Supplementary Figure 6.** Unsupervised hierarchical clustering adjacency heatmap correlation and *p*-value.

**Supplementary Figure 7:** Gene ontology and pathway analysis for the block of eight weighted gene co-expression modules associated with betaine and DMG.

**Supplementary Figure 8:** Association network on hub genes from the block of eight weighted gene co-expression modules associated with betaine and DMG.

**Supplementary Figure 9:** Heatmap of correlation between module eigengenes and cell type proportions with FDR adjusted *p*-value.

**Supplementary Figure 10:** Heatmap of correlation between sample demographic factors and nutrients and cell type proportions with FDR adjusted p-value.

### Supplementary Tables

**Supplementary Table 1:** Sample variables on RNA quality in MARBLES subjects.

**Supplementary Table 2:** Differential expression analysis on maternal gene expression and diagnosis.

**Supplementary Table 3:** ASD related significant genes from differential expression analysis.

**Supplementary Table 4:** Non-TD related significant genes from differential expression analysis.

**Supplementary Table 5:** Weighted gene co-expression network module memberships.

**Supplementary Table 6:** Weighted gene co-expression network module features, including number of transcripts and hub genes characters.

**Supplementary Table 7:** “Greenyellow” gene co-expression network module memberships on 224 transcripts.

**Supplementary Table 8:** “Greenyellow” gene co-expression network module 224 transcripts gene ontology terms and gene lists.

**Supplementary Table 9:** Eight weighted gene co-expression modules block memberships on 2,582 transcripts.

**Supplementary Table 10:** Cell type proportions in all 300 maternal peripheral blood samples estimated with CIBERSORT.

