## Supplementary_Figures for "Expression changes in immune and epigenetic gene pathways associated with nutritional metabolites in maternal blood from pregnancies resulting in autism and atypical neurodevelopment"

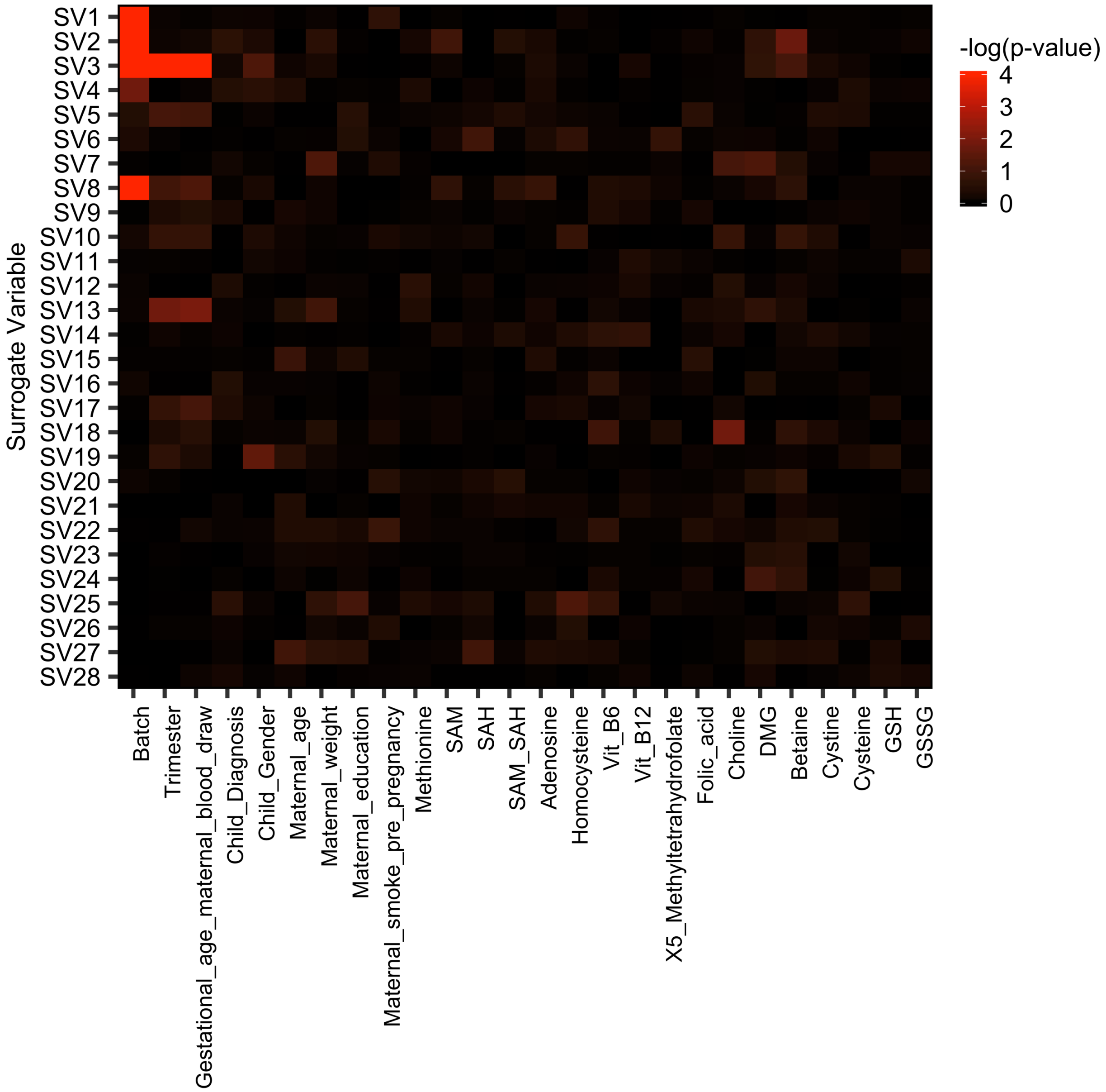


**Supplementary Figure 1.** Surrogate variable analysis in MARBLES subjects.

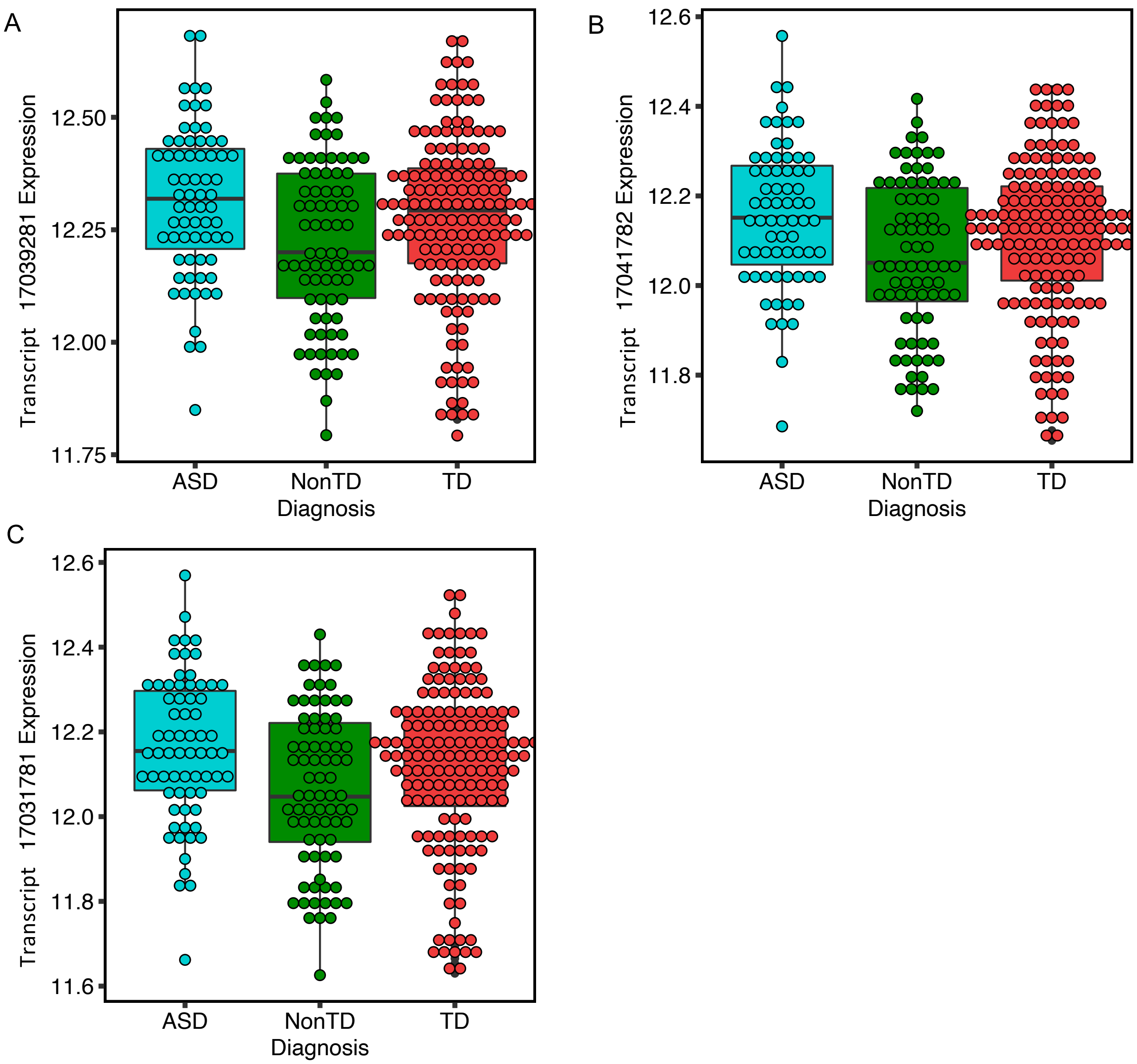


**Supplementary Figure 2.** Three transcripts at *HLA-C* reached genome-wide significance with diagnosis.

A, B and C show the expression level at each transcript. The y-axis shown the normalized and adjusted expression level. The x-axis represented three diagnosis groups, ASD, Non-TD and TD.

A) Transcript 17039281, F-test showed the significant association between expression and diagnosis (raw *p*-value = 3.16E-06, FDR adjusted *p*-value = 0.038). For ASD compared with

TD group, expression was significantly associated with ASD prior to genome-wide correction (raw *p*-value = 5.17E-04, FDR adjusted *p*-value = 0.57). In Non-TD compared to TD, expression was also significant associated with Non-TD prior to genome-wide correction (raw *p*-value = 0.017, FDR adjusted *p*-value = 0.81).

B) Transcript 17041782, diagnosis (raw *p*-value = 6.07E-06, FDR adjusted *p*-value = 0.046); ASD vs TD (raw *p*-value = 5.83E-04, FDR adjusted *p*-value = 0.57); Non-TD vs TD (raw *p*-value = 0.025, FDR adjusted *p*-value = 0.84).

C) Transcript 17031781, diagnosis (raw *p*-value = 7.96E-06, FDR adjusted *p*-value = 0.048); ASD vs TD (raw *p*-value = 6.5E-04, FDR adjusted *p*-value = 0.57); Non-TD vs TD (raw *p*-value = 0.027, FDR adjusted *p*-value = 0.86).

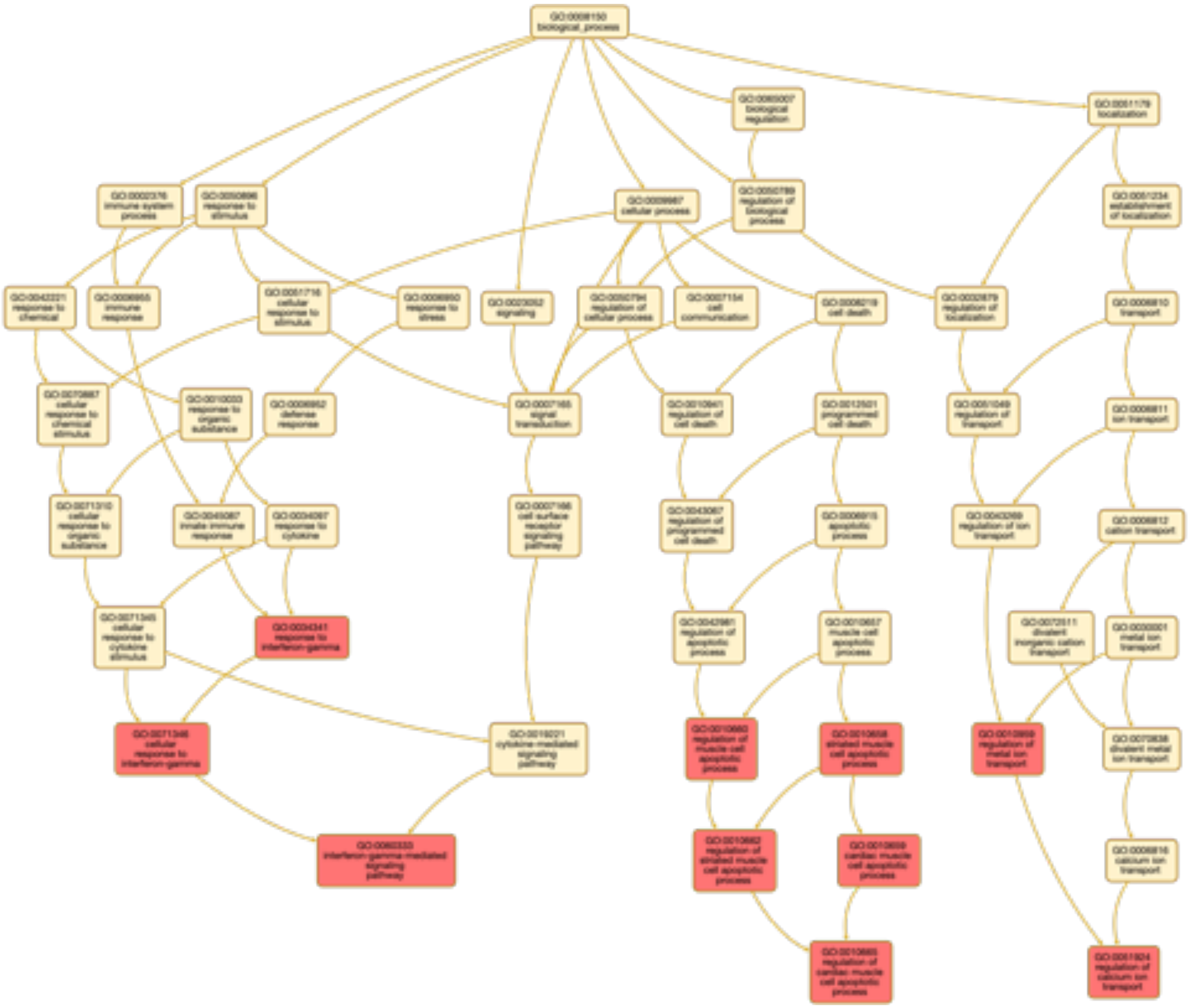

**Supplementary Figure 3.** Gene ontology and pathway analysis using directed acyclic graph (DAG) on the 218 transcripts common to ASD vs. TD and Non-TD vs. TD differentially expressed gene lists.

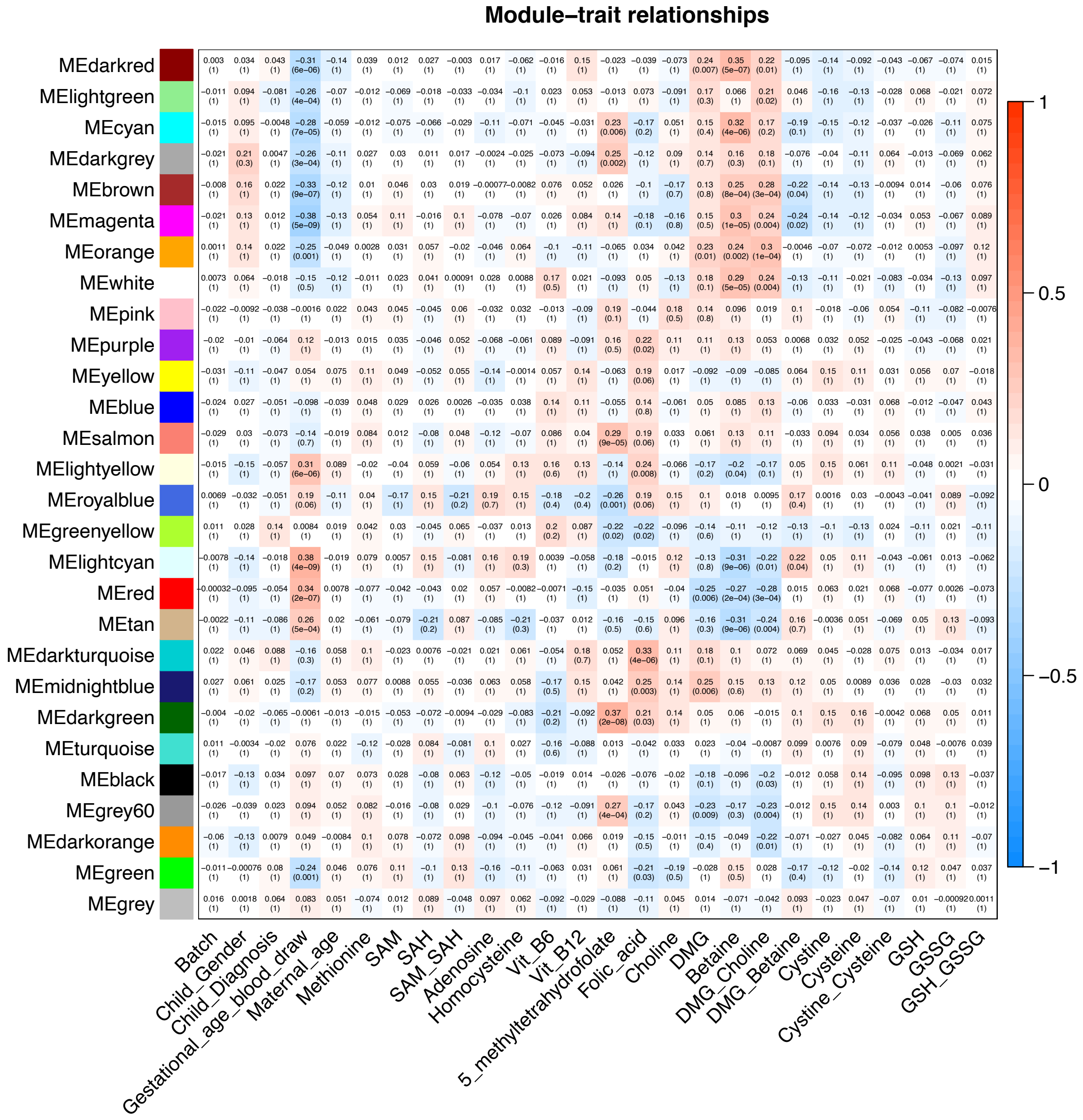


**Supplementary Figure 4.** Co-expression network modules with diagnosis, demographic factors, and maternal blood nutrient concentrations. The values in the cells represents Pearson r (adjusted *p*-value). *p*-value were adjusted for all comparisons.

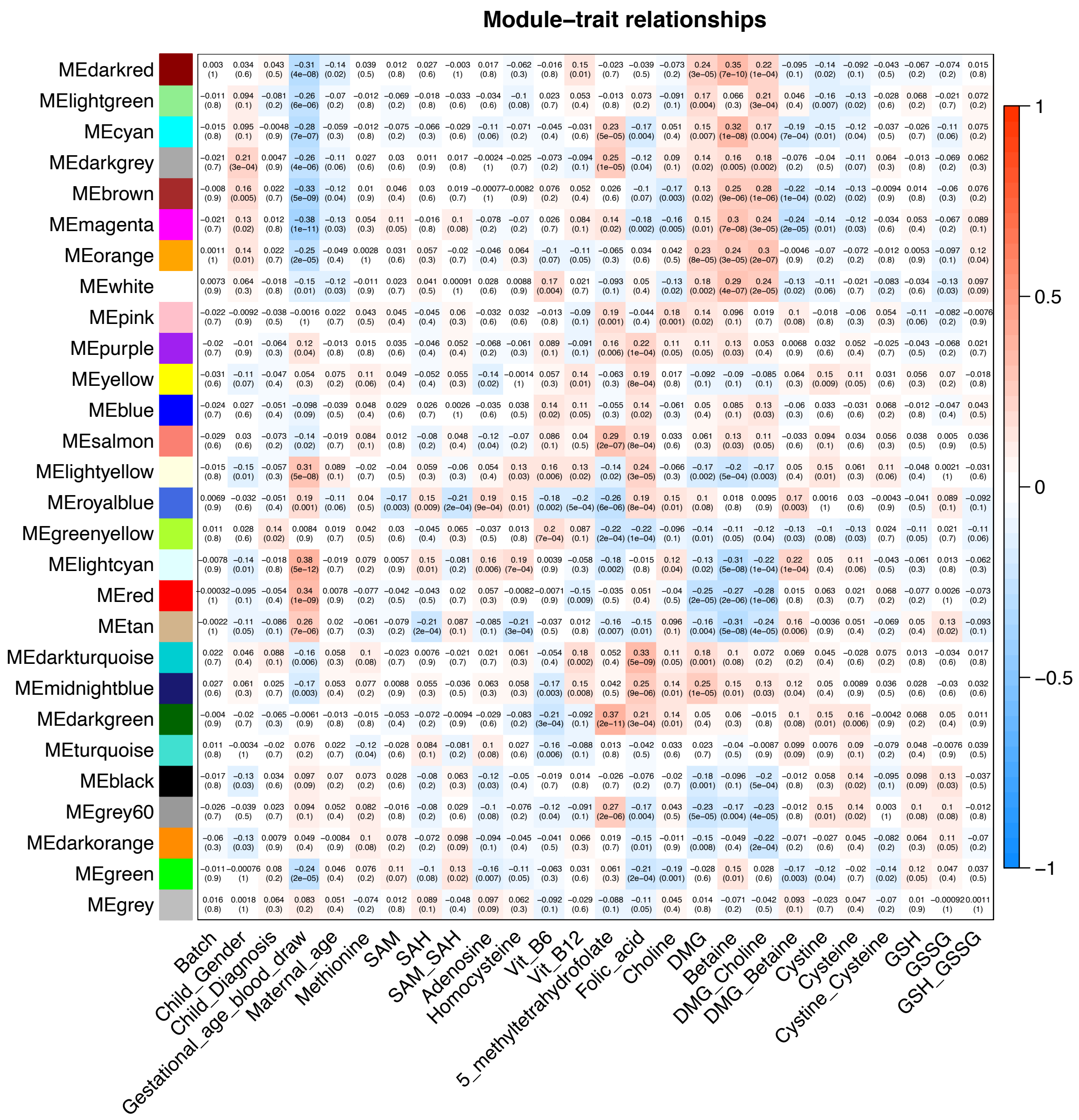


34

35 **Supplementary Figure 5.** Co-expression network modules with diagnosis, demographic

36 factors, and maternal blood nutrient concentrations. The values in the cells represents

37 Pearson r ($p$-value). $p$-value shown here were raw $p$-value without adjustment.

38

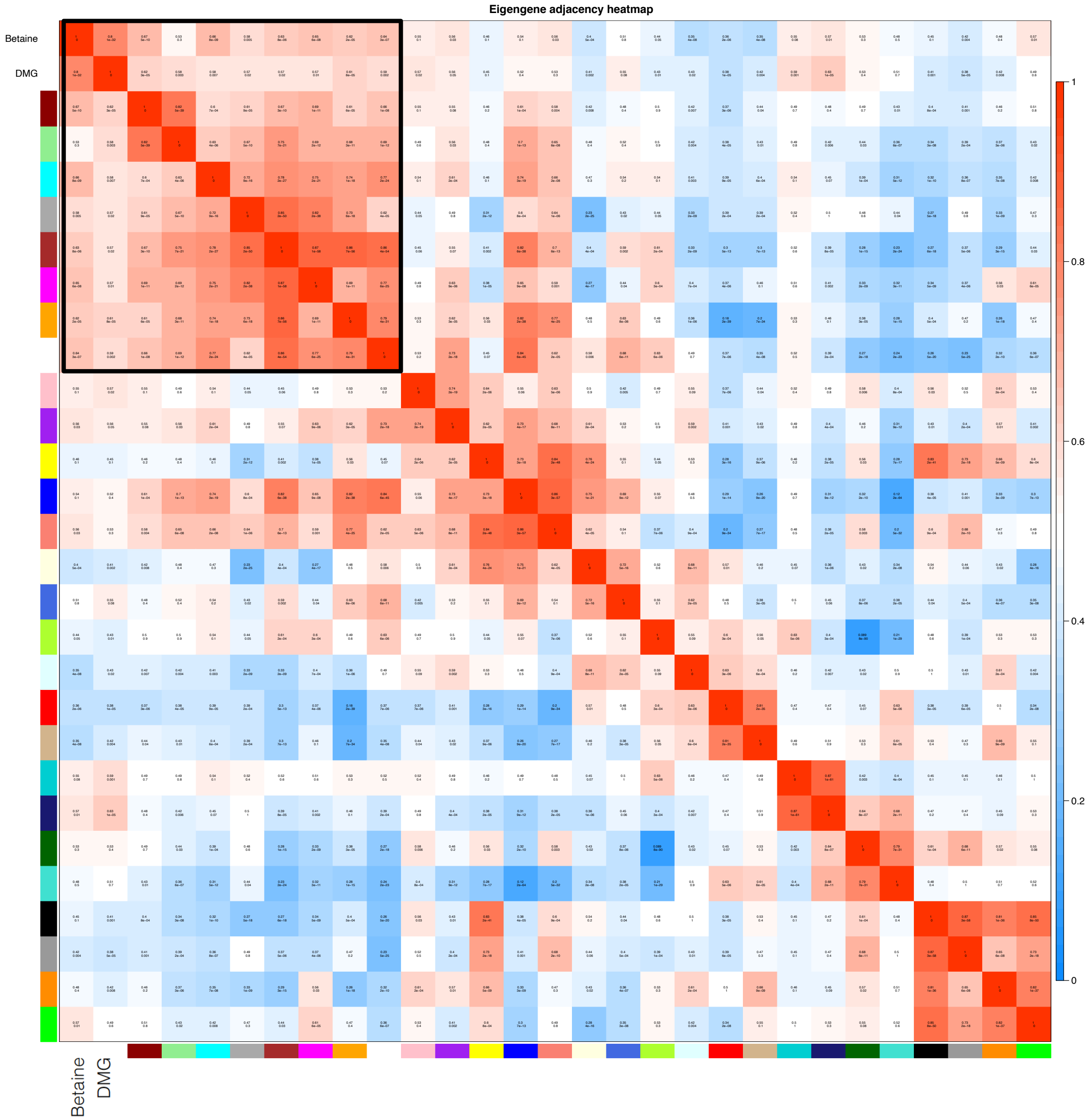


**Supplementary Figure 6.** Unsupervised hierarchical clustering adjacency heatmap correlation and *p*-value.

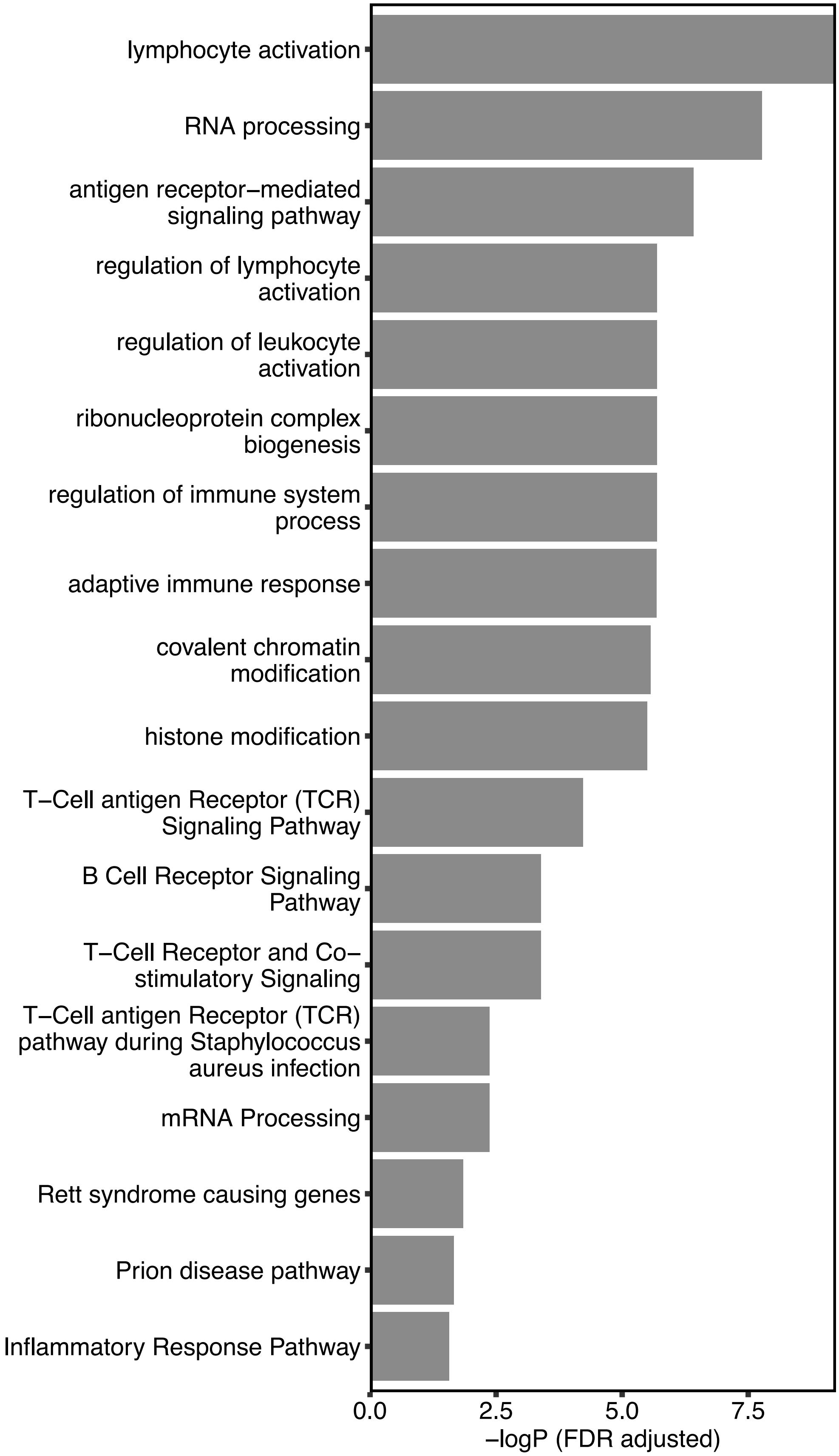
lymphocyte activation
RNA processing
antigen receptor–mediated signaling pathway
regulation of lymphocyte activation
regulation of leukocyte activation
ribonucleoprotein complex biogenesis
regulation of immune system process
adaptive immune response
covalent chromatin modification
histone modification
T–Cell antigen Receptor (TCR) Signaling Pathway
B Cell Receptor Signaling Pathway
T–Cell Receptor and Co–stimulatory Signaling
T–Cell antigen Receptor (TCR) pathway during Staphylococcus aureus infection
mRNA Processing
Rett syndrome causing genes
Prion disease pathway
Inflammatory Response Pathway
0.0
2.5
5.0
7.5
–logP (FDR adjusted)

43

**Supplementary Figure 7:** Gene ontology and pathway analysis for the block of eight weighted gene co-expression modules associated with betaine and DMG.

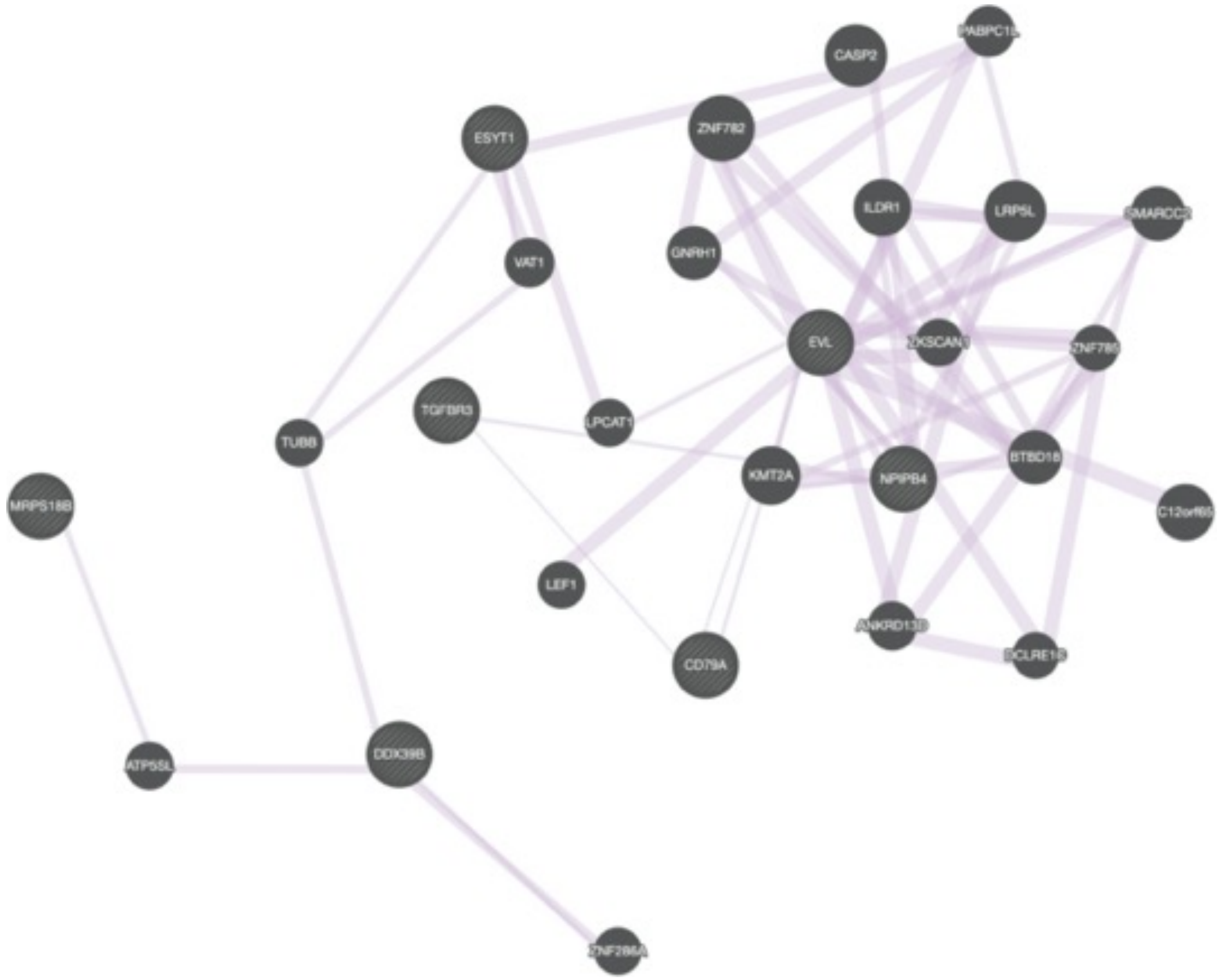


**Supplementary Figure 8:** Association network on hub genes from the block of eight weighted gene co-expression modules associated with betaine and DMG.

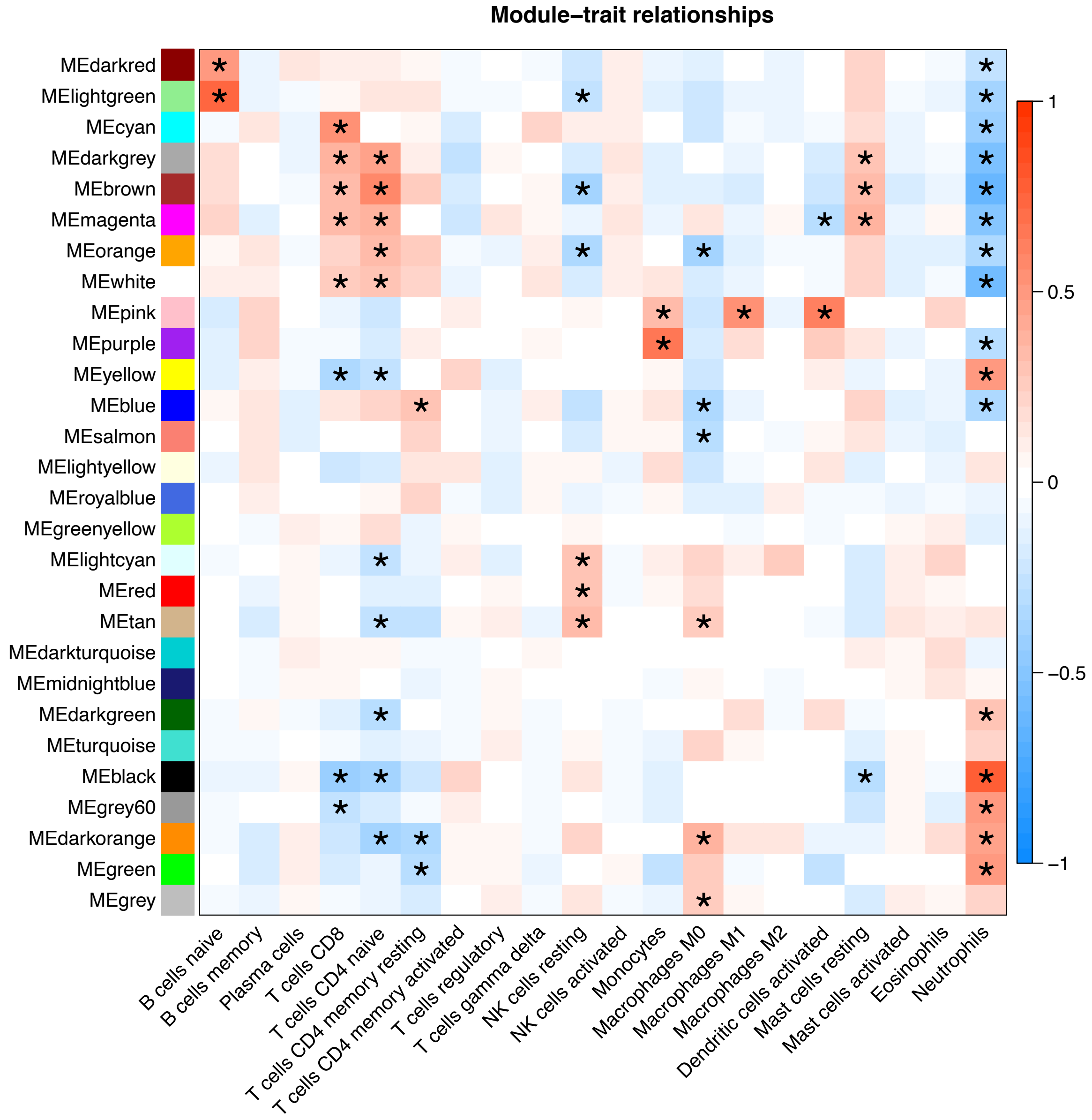


**Supplementary Figure 9:** Heatmap of correlation between module eigengenes and cell type proportions with FDR adjusted *p*-value.

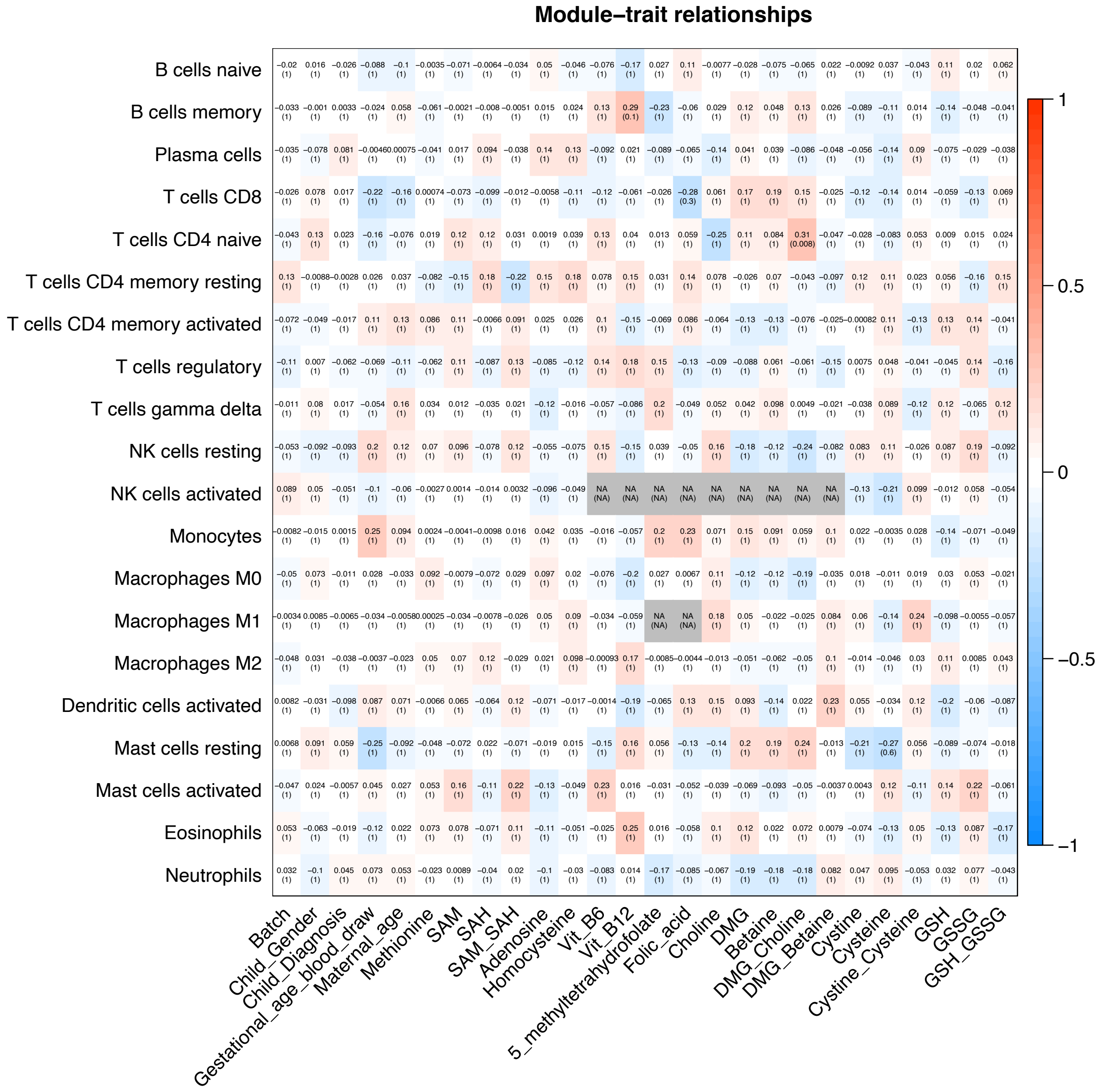


**Supplementary Figure 10:** Heatmap of correlation between sample demographic factors and nutrients and cell type proportions with FDR adjusted p-value.
